# Structure of the human Bre1 complex bound to the nucleosome

**DOI:** 10.1101/2023.03.31.535082

**Authors:** Shuhei Onishi, Kotone Uchiyama, Ko Sato, Chikako Okada, Shunsuke Kobayashi, Tomohiro Nishizawa, Osamu Nureki, Kazuhiro Ogata, Toru Sengoku

## Abstract

Histone H2B monoubiquitination (at Lys120 in humans) regulates transcription elongation and DNA repair. In humans, H2B monoubiquitination is catalyzed by the heterodimeric Bre1 complex composed of Bre1A/RNF20 and Bre1B/RNF40. The Bre1 proteins generally function as tumor suppressors, while in certain cancers, they facilitate cancer cell proliferation. To reveal the structural basis of H2BK120 ubiquitination and its regulation, we determined the cryo-EM structure of the human Bre1 complex bound to the nucleosome. The two RING domains of Bre1A and Bre1B recognize the acidic patch and the nucleosomal DNA phosphates around SHL 6.0, which are ideally located to recruit the E2 enzyme and ubiquitin for H2BK120-specific ubiquitination. Mutational experiments suggest that the two RING domains bind in both orientations and that ubiquitination occurs when Bre1A binds to the acidic patch. Our results provide insights into the relationships between Bre1 proteins and cancer and suggest that H2B monoubiquitination can be regulated by nuclesomal DNA flexibility.

## Introduction

Histones undergo various posttranslational modifications such as methylation, acetylation, and ubiquitination of lysine, which collectively regulate essentially all aspects of genome functions (*1*). These modifications induce varied biological outputs depending on their chemical types and locations. Moreover, some histone modifications are known to affect the installation, removal, and biological outputs of other modifications, known as “crosstalk” between modifications (*2*).

Ubiquitination of lysine residues is a major histone modification that regulates various genomic processes such as transcription, replication, and DNA repair (*3*). In humans, histones are monoubiquitinated at multiple lysine residues, including H3K14, H3K18, H3K23, H2AK13, H2AK15, H2AK119, and H2BK120 (corresponding to H2BK123 in yeast). Among them, monoubiquitination of H2BK120 (H2BK120ub) and H2AK119 (H2AK119ub) are involved in transcriptional regulation but play opposite roles; H2BK120ub positively regulates transcription (*4*), whereas H2AK119ub mediates transcriptional repression via the formation of facultative heterochromatin by polycomb group proteins (*5*). H2BK120ub is also important for the repair and replication of genomic DNA (*4*). Mechanistically, H2BK120ub stimulates the catalytic activity of histone H3K4 and H3K79 methyltransferases (Dot1L and MLL/Set1 family complexes, respectively) (*6-13*), thus playing a central role in a modification crosstalk that establishes a transcriptionally active chromatin environment. Moreover, H2BK120ub itself induces an accessible, open chromatin conformation (*14*) and recruits other chromatin regulators such as FACT (a histone chaperone) (*15*) and the Swi/Snf complex (a chromatin remodeling ATPase) (*16*) to regulate transcription elongation.

In yeast, H2BK123 is ubiquitinated by the Bre1 protein, which forms a heterodimer and acts as an E3 ubiquitin ligase enzyme (*17, 18*). In humans, two homologs of yeast Bre1 (Bre1A, also known as RNF20, and Bre1B, also known as RNF40) form a heterodimer responsible for H2B120 ubiquitination, with Rad6A as the E2 enzyme (*10*). The three Bre1 proteins share a C-terminal RING domain (Fig. 1a), which is required for the ubiquitination of H2B120 in humans and H2BK123 in yeasts (*17-20*). In yeast Bre1, the RING domain is minimally sufficient for nucleosomal H2BK123 ubiquitination (*21*). Its basic residues are important for the activity (*22*), presumably via interactions with the so-called “acidic patch”, a cluster of acidic residues from H2A and H2B and the hotspot of nucleosome recognition by various chromatin factors (*23*). Structural studies have demonstrated that two histone H2A-specific ubiquitin ligases, the RING1B-Bmi1 complex (ubiquitinating H2AK119) and the BRCA1-BARD1 complex (ubiquitinating H2AK125, K127, and K129), recognize the acidic patch through their basic residues in the RING domains of catalytic subunits (*24-26*). On the other hand, although the crystal structures of the RING domain of the yeast Bre1 homodimer (*27*) and human Bre1A (*28*) have been determined, no experimental structure of Bre1 proteins bound to the nucleosome has been reported.

**Fig. 1.**
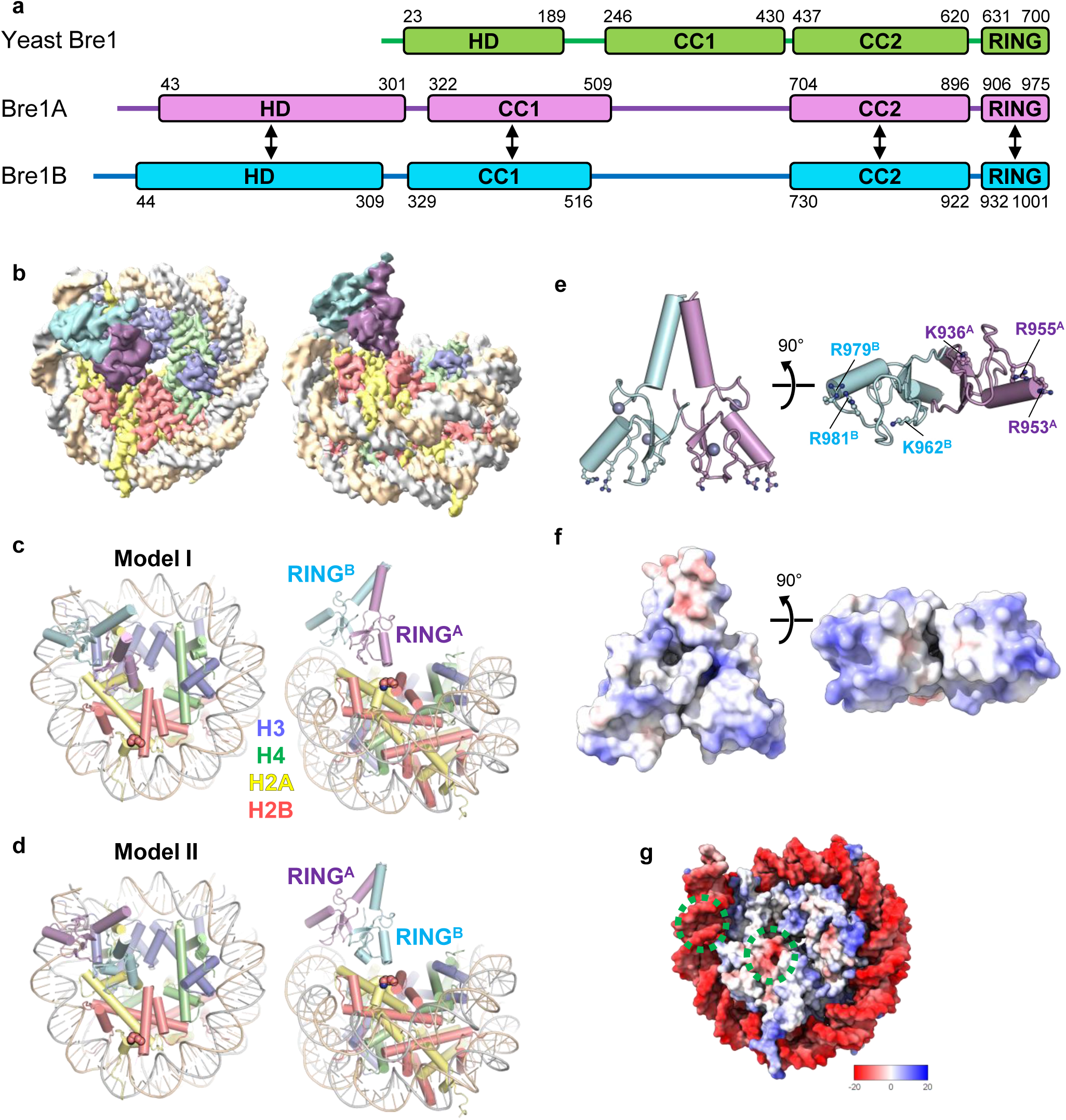
Overall structure. **a**, Domain structures of yeast Bre1 and human Bre1A and Bre1B. An N-terminal helical domain (HD) and two coiled-coil regions (CC1 and CC2) are predicted on the basis of the model structures calculated using AlphaFold. The residue numbers of the domain boundaries are shown. The arrows indicate the regions predicted to form intermolecular interactions. **b**, Cryo-EM density map in two views, colored in accordance with the model in c. **c** and **d**, Atomic models of the two RING domains of Bre1A and Bre1B bound to the nucleosome in two views. In **c** (Model I) and **d** (Model II), RING^A^ and RING^B^ bind to the acidic patch, respectively. **e**, Structure of the RING domain heterodimer (two views). Six basic residues that are important for nucleosome binding are shown. **f**, Electrostatic surface potential of the RING domain heterodimer (two views). **g**, Electrostatic surface potential of the nucleosome. The two Bre1-interaction interfaces (the acidic patch and the DNA backbone near SHL6.0) are indicated by green circles.

Accumulating evidence suggests important and complicated roles of the human Bre1 complex and H2BK120ub in cancers (*29*). In many types of cancers, expression of Bre1A or Bre1B is downregulated, and a decrease in global H2BK120ub level is associated with poor prognosis (*30-40*). Thus, Bre1A and Bre1B are generally considered tumor suppressors. On the other hand, in luminal breast cancer cells, Bre1A silencing reduces the proliferation and tumorigenic capacity of the cells, which suggests that it plays a tumor-promoting role (*31*). Bre1A expression is also required for the development of MLL-rearranged leukemia, possibly because it facilitates the transcription elongation of leukemogenic genes driven by MLL-fusion proteins (*41*). Interestingly, the knockdown of Bre1A, but not Bre1B, inhibited the proliferation of MLL-rearranged leukemia cells (*41*). Collectively, these results indicate the cancer type-specific and paralog-specific roles of human Bre1 proteins.

To reveal the structural basis of H2BK120-specific ubiquitination by Bre1 complexes and gain insights into its regulation and paralog-specific roles, we determined the cryo-EM structure of the human Bre1 complex bound to the nucleosome. The structure revealed that the RING domains of Bre1A and Bre1B directly interact with the acidic patch and the DNA phosphates of the nucleosome through their basic residues. Mutational analyses revealed that the Bre1A subunit acts as the catalytic E3 enzyme by recruiting Rad6A and identified key residues for Rad6A binding. The orientation and location of the two RING domains are consistent with the H2BK120 specificity of the Bre1 complex but differ markedly from those of the H2A-specific ubiquitin ligases. Our study provides a structural framework for the normal regulation of H2BK120 ubiquitination by the Bre1 complex and its dysregulation in pathogenic circumstances such as cancers.

## Results

### Overall structure

We reconstituted the trimeric Bre1A-Bre1B-Rad6A by mixing the coexpressed Bre1A-Bre1B heterodimer and individually expressed Rad6A and confirmed its nucleosome-binding and nucleosomal H2BK120 ubiquitination activities (Extended Data Fig. 1). Using this trimeric complex and in vitro assembled nucleosome with 147-bp DNA, we determined the cryo-EM structure of the Bre1 complex bound to the nucleosome at a global resolution of 3.1 Å. We also determined the structure of the free nucleosome at 2.74 Å resolution as a reference (Fig. 1 and Extended Data Figs. 2–4; Table 1 and Supplementary Movie 1). Although we used the full-length Bre1A-Bre1B-Rad6A heterotrimer for grid preparation, in the reconstituted cryo-EM map, only the C-terminal RING domains of Bre1A and Bre1B (hereafter designated as RING^A^ and RING^B^, respectively) were visible (Fig. 1a, 1b). This agrees with the model structures of the yeast and human Bre1 complexes calculated by AlphaFold (*42-44*), in which the C-terminal RING domains are preceded by short flexible linkers and appear to move flexibly with respect to the other regions of the complex (Extended Data Fig. 5). Previous studies have reported that the yeast and human Bre1 complexes bind Rad6 proteins via their N-terminal regions (*13, 21*), which explains why the density of Rad6A was not observed in the current EM map.

**Table 1.** Cryo-EM data acquisition, refinement, and validation statistics

|  |  |  |  |
| --- | --- | --- | --- |
| **Data collection** |  |  |  |
| Magnification | 81,000 |  |  |
| Voltage (kV) | 300 |  |  |
| Electron exposure (e<sup>-</sup>/Å<sup>2</sup>) | 58 |  |  |
| Frame | 54 |  |  |
| Defocus range (μm) | −0.8 to −1.8 |  |  |
| Pixel size (Å) | 1.05 |  |  |
| No. of micrographs | 4,653 |  |  |
| **Data processing** | Bre1 complex | Free nucleosome |  |
| EMDB ID | EMD-34274 | EMD-34275 |  |
| Symmetry imposed | C<sub>1</sub> | C<sub>1</sub> |  |
| Initial particle images | 492,704 | 492,704 |  |
| Final particle images | 22,796 | 199,682 |  |
| Map resolution (Å) | 3.1 | 2.74 |  |
| FSC threshold | 0.143 | 0.143 |  |
| **Refinement** | Model I | Model II | Free nucleosome |
| PDB ID | 8GUI | 8GUJ | 8GUK |
| Model composition |  |  |  |
| Non-hydrogen atoms | 13358 | 13269 | 12107 |
| Protein residues | 921 | 914 | 768 |
| DNA residues | 294 | 292 | 294 |
| Zinc ions | 4 | 4 | 0 |
| Model vs. data CC (mask) | 0.86 | 0.85 | 0.85 |
| Mean *B* factors (Å<sup>2</sup>) |  |  |  |
| Protein | 138.52 | 127.61 | 94.08 |
| DNA | 207.76 | 205.73 | 168.76 |
| RMS deviations |  |  |  |
| Bond lengths (Å) | 0.004 | 0.005 | 0.004 |
| Bond angles (°) | 0.548 | 0.792 | 0.720 |
| **Validation** |  |  |  |
| MolProbity score | 2.22 | 2.15 | 2.33 |
| Clash score | 12.22 | 18.50 | 14.80 |
| Ramachandran plot |  |  |  |
| Favored (%) | 96.34 | 94.30 | 96.81 |
| Allowed (%) | 3.66 | 5.48 | 3.19 |
| Outliers (%) | 0.00 | 0.22 | 0.00 |

Consistent with the previous biochemical analysis of the yeast Bre1 complex (*22*), one RING domain of the human Bre1 complex bunds to the acidic patch (Fig. 1b– d). Unexpectedly, the other RING domain binds to the DNA phosphates of one DNA strand at SHL 6.0 (Fig. 1b–d). To our knowledge, this is the first structure of any RING domain protein that directly recognizes DNA phosphates. RING^A^ and RING^B^ have positively charged molecular surfaces (Fig. 1e, 1f); thus, they bind to the negatively charged acidic patch and DNA phosphate groups (Fig. 1g) through electrostatic interactions. Bre1 binding does not induce a large conformational change in the nucleosome (Extended Data Fig. 6).

As the sequences of RING^A^ and RING^B^ are highly similar (with 86% identity; Extended Data Fig. 7), we were unable to assign the orientation of the two subunits to the density. Thus, we created two atomic models of the RING^A^-RING^B^ complex bound to the nucleosome based on the obtained density. In Model I (Fig. 1c), RING^A^ and RING^B^ bind to the acidic patch and DNA phosphates, respectively, whereas in Model II (Fig. 1d), the two RING domains have opposite orientations. On the basis of our biochemical experiments, we propose that the Bre1 complex can bind to the nucleosome in both orientations and that Model I, in which RING^A^ binds to the acidic patch, represents the catalytically active form for H2BK120 ubiquitinatio.

### Binding of the acidic patch and DNA phosphates

The detailed interactions between the acidic patch and RING^A^ or RING^B^ are shown in Fig. 2, Extended Data Fig. 8, and Supplementary Movie 2, and the density around this region is shown in Extended Data Fig. 4. Basic residues that are homologous between RING^A^ and RING^B^ (Fig. 1e and 2b and Extended Data Figs. 7 and 8) mediate the interactions with the acidic patch as follows (Bre1A and Bre1B residues are indicated with superscripted “A” and “B,”, respectively). R953^A^ and R978^B^ in Models I and II, respectively, form salt bridges with H2AE61, H2AD90, and H2AE92. These arginine residues serve as the canonical “arginine anchor” in a manner similar to that of many other chromatin factors that bind the nucleosome (*23*). R955^A^ in Model I and R981^B^ in Model II also form salt bridges with H2AE61 and H2AE64 at the position of the so-called variant arginine type 1 (*23*). In addition, the side chains of K936^A^ in Model I and K962^B^ in Model II are located at the interface with H2B, possibly contributing to nucleosome binding by forming intermolecular interactions.

**Fig. 2.**
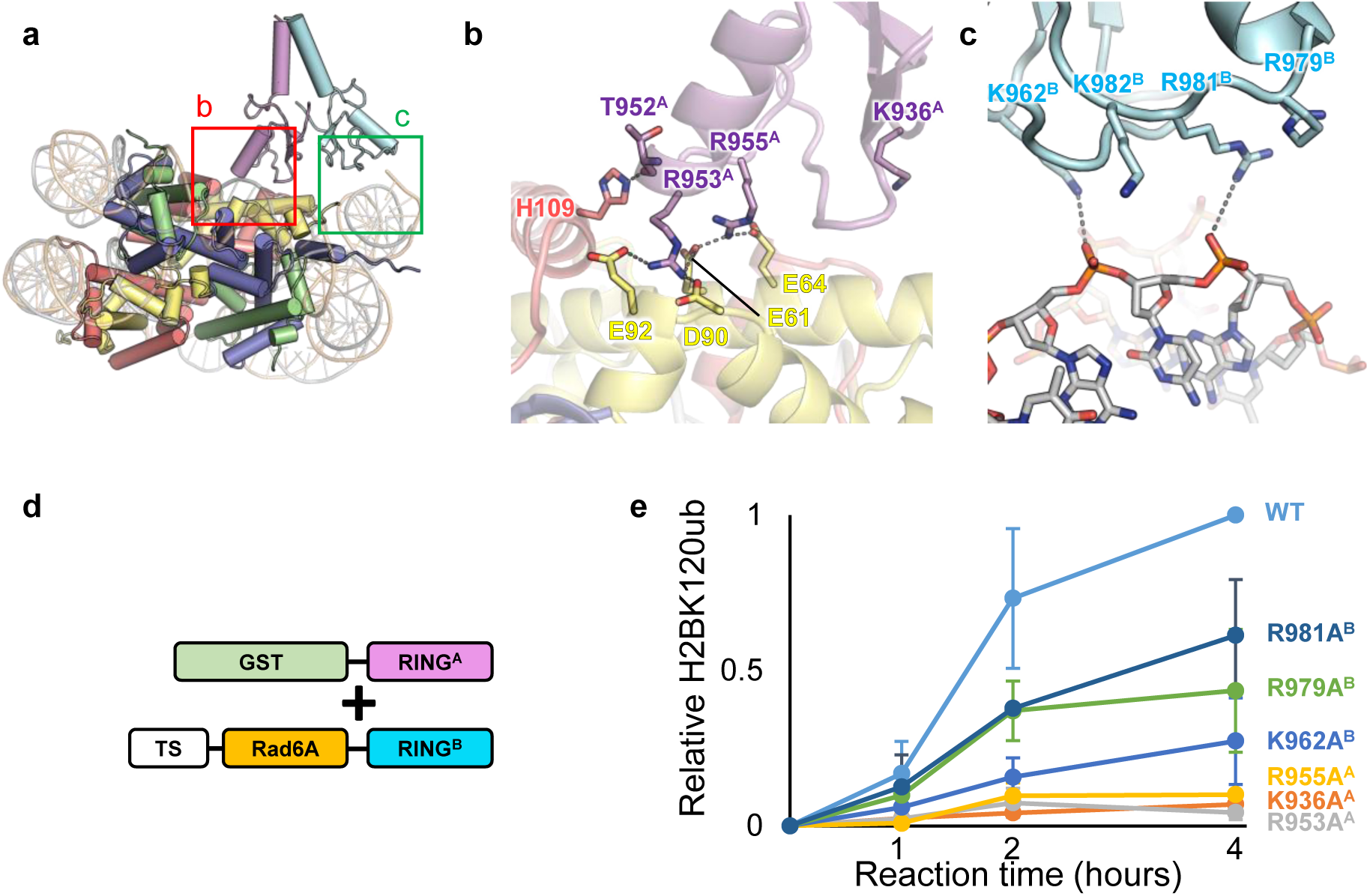
Interactions between the Bre1 complex and the nucleosome. **a**, Structure of Model I. The magnified regions in b and c are indicated. **b**, Interactions between RING^A^ and the acidic patch in Model I. Hydrophilic interactions (salt bridges and hydrogen bonds) are indicated by gray dotted lines. **c**, Basic residues of RING^B^ near the DNA phosphates in Model I. **d**, A schematic representation of the protein constructs used in the experiment shown in e. TS, Twin-Strep-tag. **e**, H2BK120 ubiquitination assay of the wild type (WT) and mutants possessing substitutions at basic residues. The signals were normalized to the WT activity at 4 hours, and the mean and standard deviation of three independent results are shown.

The other RING domain of the Bre1 complex binds the DNA phosphates around SHL 6.0 (Fig. 2a and 2c and Extended Data Fig. 8). K962^B^, R979^B^, R981^B^, and R982^B^ in Model I and K936^A^, R953^A^, R955^A^, and K956^A^ in Model II face DNA phosphates. Although the lower local resolution around this region hinders the precise modeling of their side chain atoms (Extended Data Fig. 4), these residues possibly mediate the interaction with DNA by forming salt bridges. We note that among these residues, R953^A^, R955^A^, R979^B^, and R979 ^B^ mediate interactions with the acidic patch when the Bre1 complex binds to the nucleosome in the opposite direction.

### Basic residues of the RING domains important for H2BK120 monoubiquitination

To examine the roles of the basic residues in RING^A^ and RING^B^, we prepared mutant Bre1 complexes carrying alanine mutations at the positions of K936^A^, R953^A^, R955^A^, K962^B^, R979^B^, and R981^B^. To focus on the functional role of the RING domains and facilitate enzymatic analysis, we created constructs expressing RING^A^ or RING^B^ fused to Twin-Strep-tag and Rad6A and coexpressed each of them with the GST-fused RING domain of the other Bre1 protein (Fig. 2d and Extended Data Fig. 9a). A similar fusion strategy was used to analyze the activity of the yeast Bre1 complex (*21, 22*). To purify the heterodimeric complexes, the coexpressed proteins were purified by serial affinity chromatography steps using Strep-Tactin and glutathione resins and then used for enzymatic analysis.

The results of the enzymatic analysis using GST-RING^A^ and Twin-Strep-Rad6A-RING^B^ are shown in Fig. 2e, whereas those of the enzymatic analysis using GST-RING^B^and Twin-Strep-Rad6A-RING^A^ are shown in Extended Data Fig. 9b. All six mutations showed reduced activity, albeit at varying degrees. The K936A^A^, R953A^A^, and R955A^A^ mutations almost completely abolished the activity in both combinations. The K962A^B^, R979A^B^, and R981A^B^ mutations showed milder effects but still showed significantly reduced activity when GST-RING^A^ and Twin-Strep-Rad6A-RING^B^ were used, while they greatly reduced the activity when GST-RING^B^ and Twin-Strep-Rad6A-RING^A^ were used. These results demonstrate the importance of the interaction between the basic residues of the two RING domains and the acidic patch and nucleosomal DNA observed in the current structure for H2BK120 ubiquitination.

### Bre1A as a catalytic subunit

The mutational analysis results showed that mutations on Bre1B had milder effects, which suggests that Bre1A and Bre1B play asymmetrical functions in catalysis, even though their sequences are similar. Consistent with this, a previous study showed that the ectopic expression and knockdown of Bre1A, not Bre1B, affected the H2BK120ub level in HEK293T cells (*19*). Thus, we addressed whether the RING domains of Bre1A and Bre1B may perform asymmetrical functions. Many RING-type E3 enzymes, including the H2A-specific enzymes RING1B and BRCA1, are known to bind non-catalytic RING proteins to form RING domain heterodimers, in which the catalytic RING subunit recruits the E2 enzyme and ubiquitin for site-specific ubiquitination (*45*). To examine if Bre1 proteins can asymmetrically interact with its cognate E2 protein, Rad6A, we first created model structures of Rad6A bound to the Bre1 RING domains by superposing with the structures of a known RING E3-E2-ubiquitin complex, the RNF4-UbcH5A ubiquitin complex (*46*) (PDB ID 4AP4), followed by the superposition of human Rad6A (*47*) (PDB ID 6CYO) on UbcH5A (Fig. 3a, Supplementary Movie 1). Previous reports have created model structures of yeast and human Bre1 complexes with bound Rad6 in a similar manner, which were supported by cross-linking and mass spectrometry analyses or a mutational assay (*22, 28*). Our human Bre1-Rad6A model suggests that if RING^A^ binds Rad6, the interaction may involve Bre1A residues C924^A^, N926^A^, M927^A^, T948^A^, R949^A^, T952^A^, Q954^A^, K956^A^, P958^A^, and K959^A^. Bre1A and Bre1B have different amino acids only at three (M927^A^/T953^B^, T948^A^/G974^B^, and T952^A^/A978^B^) of these positions (Fig. 3b, Extended Data Fig. 7). Furthermore, in the model structure, the side chains of T948^A^ and T952^A^ are located within the hydrogen-bonding distance of the N65 and K66 side chains of Rad6A (Fig. 3c). These observations suggest that the two positions, T948^A^/G974^B^ and T952^A^/A978^B^, are likely located at the interaction interface with Rad6A and that the amino acids at these positions may be involved in the asymmetrical functions of Bre1A and Bre1B.

**Fig. 3.**
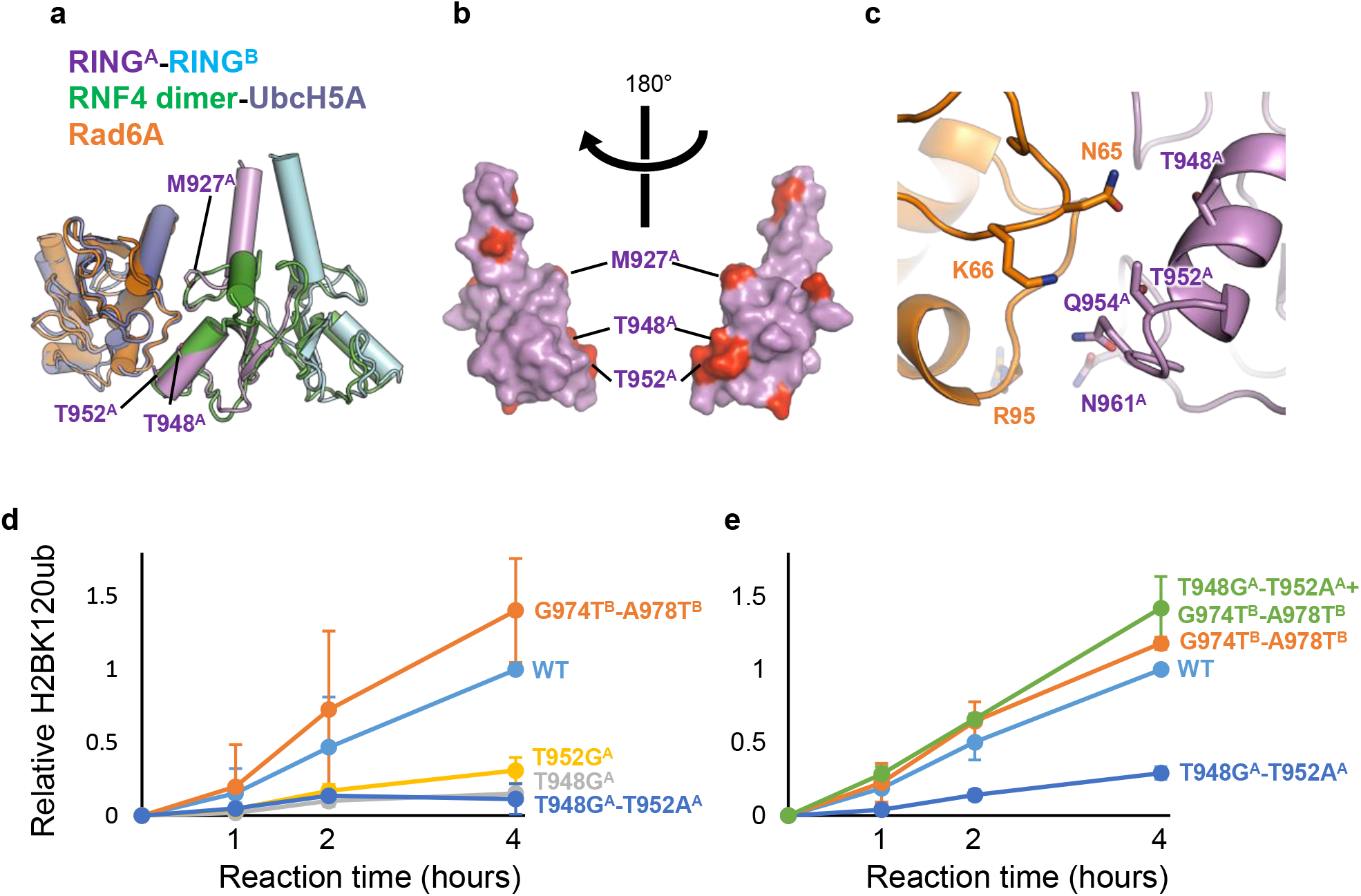
Identification of Bre1A residues likely to mediate Rad6A binding. **a**, Structural modeling of the RING^A^-RING^B^-Rad6A complex. The current structure of the RING^A^-RING^B^ complex is superposed on that of the RNF4 dimer bound to UbcH5A and ubiquitin (PDB ID 4AP4, ubiquitin not shown), and then the structure of Rad6A (PDB ID 6CYO) is further superposed on that of UbcH5A. **b**, RING^A^ residues that are not conserved in RING^B^. The RING^A^ structure is shown as a surface representation, and 10 non-conserved residues are colored red. Three residues near a possible Rad6A binding region are labeled. **c**, Close-up view of the RING^A^- RING^B^-Rad6A model in b. **d** and **e,** H2BK120 ubiquitination assay of the wild type (WT) and mutants possessing substitutions at the possible Rad6A binding region. The constructs shown in Fig. 2d and their mutants were used.

To directly test whether T948^A^/G974^B^ and T952^A^/A978^B^ play important roles, we purified Bre1-Rad6 fusion complexes with the Bre1A subunit carrying a single mutation or double Bre1B-like mutations as follows: T948G^A^, T952A^A^, or 948G^A^-T952G^A^. We then analyzed their nucleosomal H2BK120 ubiquitination activity. As shown in Fig. 3d, the T948G^A^, T952A^A^, and T948G^A^-T952G^A^ mutations all significantly reduced the activity of the complexes. To further validate the importance of threonine at these positions, we created two Bre1 complexes, one containing the Bre1B subunit carrying Bre1A-like dual mutations (G974T^B^-A978T^B^) and the wild-type Bre1A subunit, and the other carrying Bre1B-like Bre1A and Bre1A-like Bre1 subunits (Bre1A T948G^A^-T952G^A^ + Bre1B G974T^B^-A978T^B^). As shown in Fig. 3e, the complex carrying the G974T^B^-A978T^B^ mutations exhibited slightly higher activity than the wild-type complex, and the activity of the complex carrying T948G^A^-T952G^A^ and G974T^B^-A978T^B^ was comparable with that of the wild-type complex. These results demonstrate that Bre1A functions as a catalytic subunit to recruit the E2-ubiqution complex for H2BK120 mono ubiquitination and that both T948A and T952A are important for Rad6A binding.

### A model for H2BK120-specific ubiquitination

Fig. 4a shows the model structure of the RING^A^-RING^B^-Rad6A ubiquitin bound to the nucleosome, which is based on Model I, in which RING^A^ binds the acidic patch. In this model, the lysine residue closest to ubiquitin Gly76 is H2BK120 (Fig. 4b, Supplementary Movie 3), and the distance between the Cα atom of H2BK120 and the carbonyl carbon of ubiquitin G76 is 8.4 Å. The second closest lysine is H2BK116, whose Cα atom is located 10.5 Å away from the carbonyl carbon of ubiquitin G76 and might be too farther away for ubiquitin transfer (Fig. 4b). Unlike the C-terminal ubiquitination sites of H2A, which are structurally flexible, H2BK117 and H2BK120 are located on the C-terminal α-helix of H2B. The helical structure should exhibit limited backbone flexibility, which may hinder the access of H2BK117 to the ubiquitination catalytic center. Taken together, our model structure of the RING^A^-RING^B^-Rad6A-ubiquitin complex bound to the nucleosome is consistent with the H2BK120 specificity of the human Bre1 complex.

**Fig. 4.**
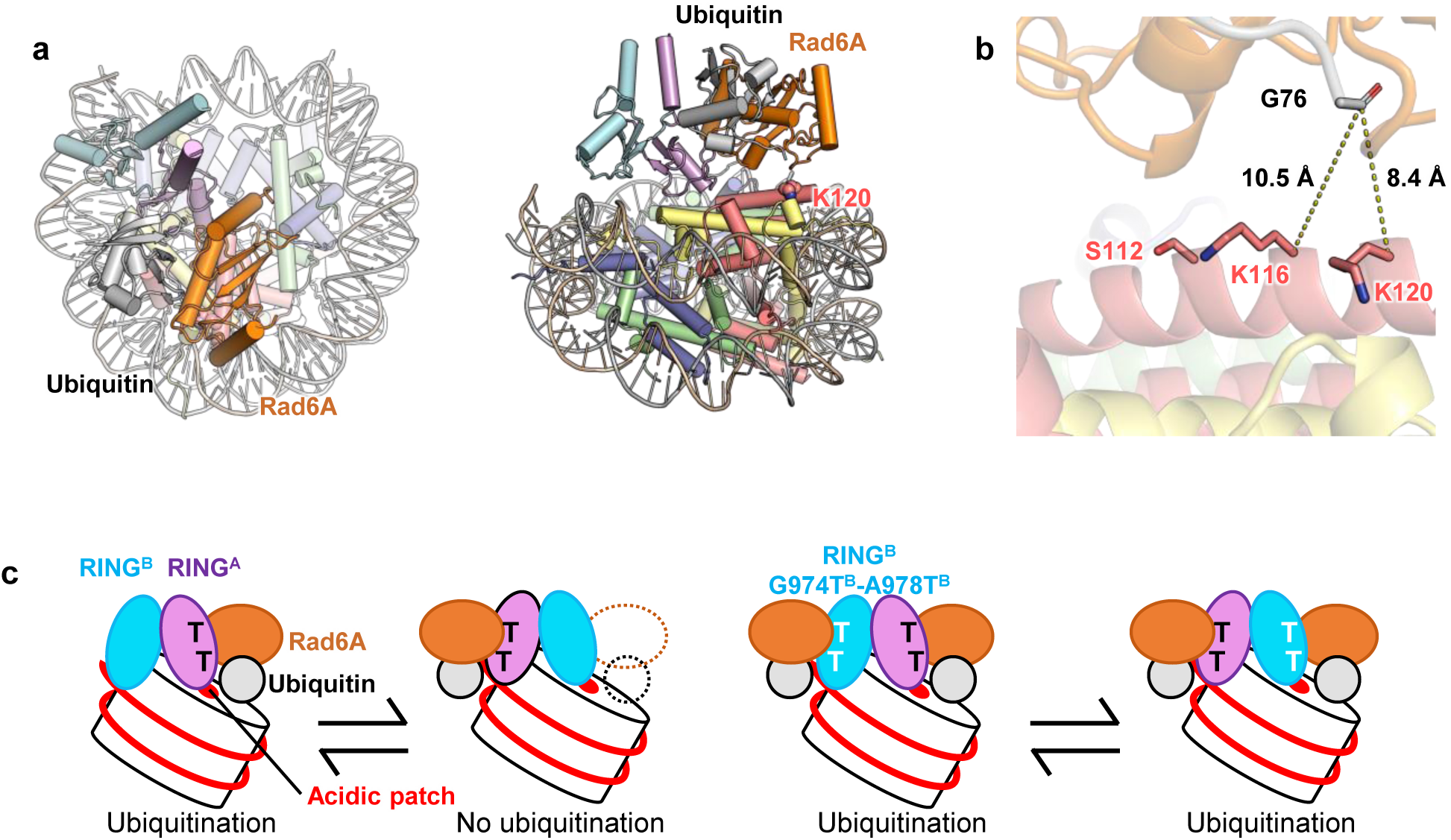
A model for H2BK120-specific ubiquitination by the Bre1 complex. **a**, Model structure of the RINGA-RINGB-Rad6A-ubiquitin complex bound to the nucleosome in two views. **b**, Close-up view of ubiquitin and H2B. Two lysine residues (H2BK120 and H2BK116) near G76 of ubiquitin are shown. H2BS112, whose GlcNAcylation stimulates H2BK120 ubiquitination, is also shown. **c**, Proposed mechanistic model. The wild-type Bre1 complex can bind to the nucleosome in two orientations, but H2BK120 ubiquitination occurs only when Bre1A binds to the acidic patch, as RING^A^, but not RING^B^, can recruit Rad6A and ubiquitin. Bre1B with G974T^B^-A978T^B^ double substitution can recruit Rad6A and ubiquitin; thus, H2BK120 ubiquitination occurs in both binding modes.

Our structural model also suggests that the catalytic RING domain should bind to the acidic patch to position the E2 enzyme and ubiquitin near H2BK120 for ubiquitination, as the other RING domain contacting the DNA phosphates is farther away from H2BK120 and faces the opposite direction. Thus, when the wild-type Bre1 complex ubiquitinates H2BK120, the catalytically active RING^A^ should bind to the acidic patch. Meanwhile, the results of our mutational experiments also showed that when Bre1A-type mutations (G974T^B^ and A978T^B^) were introduced, RING^B^ could function as the catalytic E3 enzyme. This suggests that RING^B^ also binds to the acidic patch, which agrees with the high sequence similarity between RING^A^ and RING^B^ and the perfect conservation of the two arginine residues that binds to the acidic patch (R953^A^/R979^B^ and R955^A^/R981^B^).

On the basis of these observations, we propose a mechanistic model of how the human Bre1 complex ubiquitinates H2BK120 (Fig. 4c). The Bre1 complex interacts with the nucleosome in at least two regions: the acidic patch and the DNA phosphates around SHL 6.0. The two regions are recognized on the basis of the basic surfaces of the two RING domains. The Bre1 complex interacts with the nucleosome in two possible orientations of their RING domains, and H2BK120 ubiquitination occurs only when RING^A^ binds to the acidic patch. R953^A^ and R955^A^ are the canonical arginine anchor and variant arginine type 1, respectively, for recognizing the acidic patch and are thus essential for H2BK120 ubiquitination. On the other hand, basic residues in RING^B^, such as K962^B^, R979^B^, and R981^B^, have a rather supportive role in orienting the catalytic RING^A^ domain and providing additional affinity for the nucleosome via interaction with the nucleosomal DNA phosphates. When Bre1A-type mutations (G974T^B^ and/or A978T^B^) are introduced, the mutant Bre1 complex can catalyze H2BK120 ubiquitination in both orientations, as both RING domains can recruit Rad6A and ubiquitin.

### Comparison with H2A ubiquitin ligases

The structural comparison between the current Bre1 complex (Fig. 5a) and two other histone ubiquitin ligases recognizing the nucleosome (the RING1B-Bmi1 complex ubiquitinating H2AK119, shown in Fig. 5b, and the BRCA1-BARD1 complex ubiquitinating H2AK125, K127, and K129, shown in Fig. 5c) revealed a clear similarity and difference between their nucleosome recognition mechanisms. In all structures, the catalytic RING domain recognizes the acidic patch using the conserved arginine residues as the arginine anchor, clarifying a common mechanism adopted by all structurally characterized histone ubiquitin ligases that recognize the nucleosome. On the other hand, the orientation of the catalytic RING domain in the Bre1 complex differs substantially from that of the two H2A ubiquitin ligases, consistent with their different residue specificity. Moreover, the non-catalytic RING domains interact with the nucleosome differently. Unlike the Bre1 complex, whose non-catalytic RING domain (RING^B^) binds to DNA phosphates, the non-catalytic RING domains of the two H2A ubiquitin ligases (Bmi1 and BARD1) do not possess the basic surface and thus do not bind to DNA. Instead, Bmi1 contacts residues 76–79 of histone H3, whereas BARD1 interacts mainly with the H2B/H4 cleft, resulting in distinct residue specificities on the H2A C-terminal region. In summary, the two H2A ubiquitin ligases bind to nucleosomes in similar but distinct ways to specifically ubiquitinate different H2A lysine residues, whereas the Bre1 complex binds in a nearly opposite direction for H2BK120-specific ubiquitination.

**Fig. 5.**
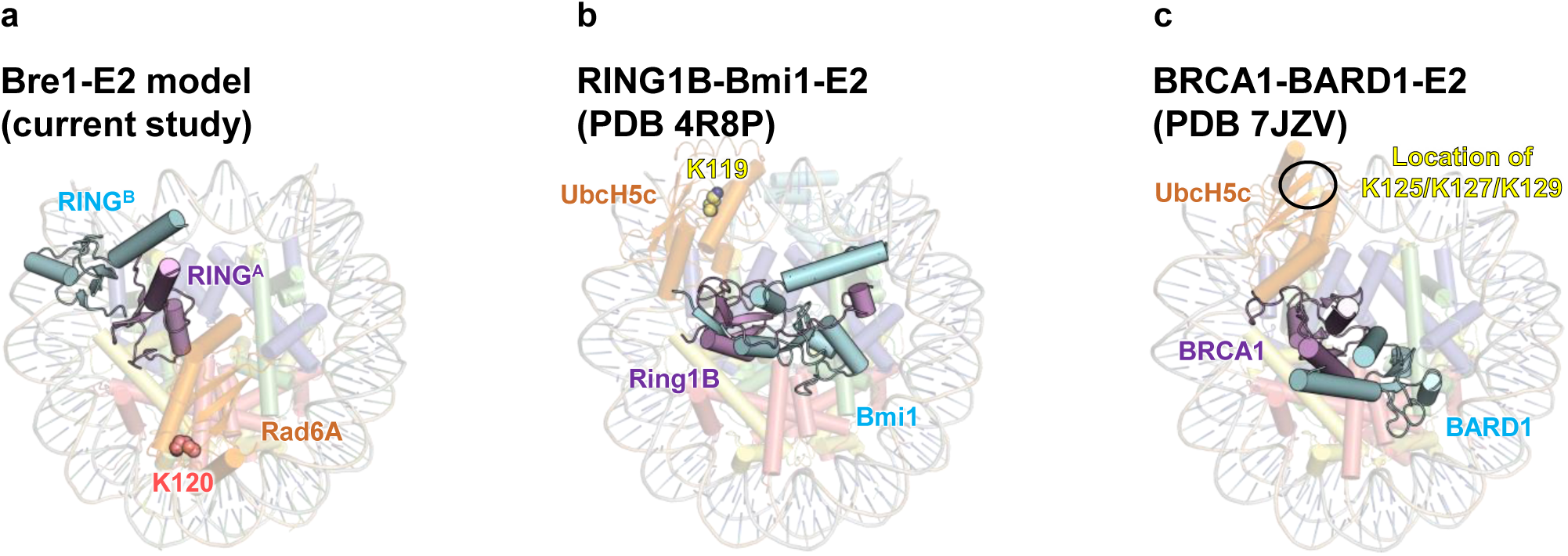
Comparison of nucleosomal histone ubiquitin ligases. **a**, Model structure of RING^A^- RING^B^-Rad6A-ubiquitin bound with the nucleosome (ubiquitin omitted for clarity). **b**, Structure of the RING1B-Bmi1 complex and its E2 enzyme, UbcH5c, bound to the nucleosome. **c**, Structure of the BRCA1-BARD1 complex and its E2 enzyme, UbcH5c, bound to the nucleosome. The target residues (H2AK125, K127, and K129) are disordered in the structure.

## Discussion

Our structural and mutational analyses revealed that the human Bre1 complex recognizes the acidic patch and DNA phosphates for H2BK120-specific ubiquitination and identified Bre1A as the catalytic subunit that recruits the cognate E2 enzyme (Rad6A) and ubiquitin. These findings rationalize previous biochemical data about yeast and human Bre1 complexes, reconcile the apparently contradictory roles of Bre1A and Bre1B in cancer development, and provide a structural framework for understanding the cellular mechanisms regulating H2BK120 ubiquitination and developing potential therapeutics.

Previous studies have examined how H2BK120 ubiquitination is affected by other histone modifications. Histones are *O*-GlcNAcylated at multiple residues by *O*-linked *N*-acetylglucosamine transferase (*48*), and GlcNAcylation at H2BS112 was shown to facilitate H2BK120 ubiquitination by the Bre1 complex in vitro and in vivo (*49*). A systematic study using semi-synthesized nucleosomes confirmed this facilitating effect by H2BS112 GlcNAcylation (*50*). In our model structure of RING^A^-RING^B^-Rad6A-ubiquitin bound to the nucleosome, H2BS112 is located close to Rad6A (Fig. 5b). It is possible that H2BS112GlcNAc facilitates H2BK120 ubiquitination by interacting with Rad6, thereby increasing the binding affinity or correctly positioning Rad6A and ubiquitin suitable for ubiquitination. The systematic study using semi-synthesized nucleosomes also revealed multiple histone H3 residues whose modification or mutation stimulates H2BK120 ubiquitination (*50*). These modifications and mutations, including H3Y41ph and H3K56ac, cluster near the DNA entry/exit site and tend to weaken histone-DNA contacts just around the Bre1-bound site (Extended Data Fig. 10). A possible explanation is that these histone-DNA contacts weakened by the modifications affect the flexibility of the DNA around SHL 6.0, which may facilitate the productive binding of the Bre1 complex on the DNA. Further studies are needed to test this hypothesis and to examine whether such a regulatory mechanism may operate in a physiologically significant manner.

Although Bre1 proteins are generally considered tumor suppressors, various reports have suggested that their roles in cancers are complicated and context dependent. Several cancer cells, such as MLL-rearranged leukemia cells (*41*) and luminal breast cancer cells (*31*), require Ber1A for proliferation. High levels of H2BK120 and Bre1 gene expressions are associated with poor survival in some cancers (*31, 51*). These results suggest that the Bre1 complex plays a tumor-promoting role in these cancers. Our structural and biochemical analyses pave the way for the rational design of Bre1 inhibitors as potential therapeutics for such cancers. Moreover, the identification of Bre1A as the catalytic E3 subunit is consistent with the finding of a study that showed that Bre1A, but not Bre1B, is required for the development of MLL-rearranged leukemia (*41*).

## Online Methods

### AlphaFold calculation

The model structures of the full-length yeast Bre1 homodimer and human Bre1 heterodimer were calculated using the program LocalColabFold (*44*), with AlphaFold2 (*42*) as the calculation engine.

### Protein and DNA sequences

The protein and nucleosomal DNA sequences used in this study are shown in Supplementary Table 1.

### Purification of the Bre1A-Bre1B-Rad6A complex

The full-length human Rad6A sequence was cloned into a modified pET32 plasmid (Roche) and expressed as a fusion protein with the N-terminal thioredoxin-His6-SUMOstar tag. The plasmid was introduced in Rosetta2(DE3) pLysS cells, and the protein expression was induced by adding 0.2 mM isopropyl-β-D-thiogalactoside (IPTG) and culturing overnight at 20°C. The harvested cells were suspended in 20 mM Na phosphate at pH 7.3, 500 mM NaCl, 1 mM DTT, 0.1 mM phenylmethylsulfonyl fluoride (PMSF), and 1% Triton X-100 and disrupted by sonication. Clarified lysate was loaded onto a cOmplete His-Tag column (Roche). The protein was eluted with 20 mM Na phosphate at pH 7.3, 500 mM NaCl, 1 mM DTT, and 500 mM imidazole. For the reconstitution of the Bre1A-Bre1B-Rad6A complex, the tag-fused Rad6A was flash-frozen in liquid nitrogen and stored at −80°C. For the histone H2B ubiquitination assay, the tag-fused Rad6A was mixed with SUMOstar protease and dialyzed against 20 mM Tris-HCl at pH 7.5, 150 mM NaCl, and 1 mM DTT overnight. The digested sample was then mixed with an equal amount of water and loaded onto tandemly connected HisTrap HP and HiTrap Q columns (Cytiva) equilibrated with 20 mM Tris-HCl at pH 7.5, 100 mM NaCl, 20 mM imidazole, and 1 mM DTT. After disconnecting the HisTrap column, the bound Rad6A protein was eluted from the HiTrapQ column with a linear gradient from 100 mM to 1 M of NaCl in 20 mM Tris-HCl at pH 7.5, 20 mM imidazole, and 1 mM DTT. The fractions containing Rad6A were pooled, concentrated, buffer exchanged to 10 mM HEPES-Na at pH 7.5, 150 mM NaCl, 1 mM DTT, flash-frozen with liquid nitrogen, and stored at −80°C.

For the structural study, the full-length sequences of human Bre1A and Bre1B were first individually cloned into pACEBac1 (Geneva Biotech) with the N-terminal His6-SUMOstar tag and the N-terminal Twin-Strep-His6-SUMOstar tag, respectively. The two expression cassettes were then integrated into a single pACEBac1 plasmid to construct a bicistronic vector expressing both proteins. Baculovirus production in Sf9 cells was conducted using the Bac-to-Bac system (ThermoFisher), in accordance with the manufacturer’s instructions. The full-length Bre1A-Bre1B complex was expressed using the bicistronic baculovirus in High Five cells, which were collected 48 h after viral infection. The cell pellets were suspended in 20 mM Tris-HCl at pH 8.0, 200 mM NaCl, 1 mM DTT, 0.1 m PMSF, 0.5% Triton X-100, and cOmplete ULTRA protease inhibitor cocktail (Roche) and disrupted by sonication. Clarified lysate was loaded onto a Strep-Trap HP column (Cytiva), and the complex was eluted in a buffer containing 20 mM Tris-HCl at pH 8.0, 200 mM NaCl, 1 mM DTT, and 2.5 mM desthiobiotin. To reconstitute the trimeric Bre1A-Bre1B-Rad6A complex, the eluted fractions containing the tagged Bre1A-Bre1B complex were mixed with the tag-fused Rad6A and SUMOstar protease and incubated at 4°C overnight. The tag-removed, reconstituted ternary complex was then loaded onto a HiTrap Heparin column (Cytiva) equilibrated with 20 mM Tris-HCl at pH 7.5, 100 mM NaCl, and 1 mM DTT, and eluted with a linear gradient from 100 mM to 1 M NaCl in 20 mM Tris-HCl at pH 7.5 and 1 mM DTT. The complex was further purified with a Superose 6 column (Cytiva) equilibrated with 20 mM Hepes-Na at pH 7.5, 200 mM NaCl, and 1 mM DTT, concentrated, flash-frozen in liquid nitrogen, and stored at −80°C.

The DNA sequences encoding the RING domain of Bre1A or Bre1B were cloned into pGEX-6P-1 (Cytiva) to express GST-RING^A^or GST-RING^B^ and cloned into a modified pET plasmid (Merck) to express Twin-Strep-Rad6A-RING^A^ or Twin-Strep-Rad6A-RING^B^. To prepare the RING domain heterodimers, two plasmids (pGEX-6P-RING^A^ and pET-Twin-Strep-Rad6A-RING^B^ or pGEX-6P-RING^B^ and pET-Twin-Strep-Rad6A-RING^A^) were simultaneously introduced into Rosetta2(DE3)pLysS cells. Protein expression was induced with 0.2 mM IPTG and overnight culture at 16°C. The harvested cells were suspended in 40 mM K_2_HPO_4_, 10 mM KH_2_PO_4_, 500 mM NaCl, 1 mM DTT, 0.1 mM PMSF, and 0.5% Triton X-100 and lysed by sonication. The clarified lysate was applied to a StrepTrap column (Cytiva), and the bound heterodimers were eluted with 50 mM Tris-HCl at pH 7.5, 500 mM NaCl, 1 mM DTT, 2.5 mM desthiobiotin. The eluents were further purified with a GSTrap column (cytiva) and eluted with 50 mM Tris-HCl at pH 8.0, 500 mM NaCl, and 10 mM reduced glutathione. The purified heterodimers were flash-frozen in liquid nitrogen and stored at −80°C.

The plasmids encoding GST- and His-tagged human E1 (pGEX-6P-HsE1-His8) and ubiquitin (pET26b-Ub) were kind gifts from Dr. Yusuke Sato (Tottori University). pGEX-6P-HsE1-His8 was introduced into Rosetta2(DE3)pLysS cells, and protein expression was induced by adding 0.3 mM IPTG and culturing overnight at 20°C. The harvested cells were suspended in 40 mM K_2_HPO_4_, 10 mM KH_2_PO_4_, 20 mM imidazole, 500 mM NaCl, 1 mM DTT, 0.1 mM PMSF, and 0.5% Triton X-100 and lysed by sonication. The clarified lysate was applied to a HisTrap column, and the protein was eluted with a linear gradient from 20 mM to 500 mM of imidazole in 40 mM K_2_HPO_4_, 10 mM KH_2_PO_4_, 500 mM NaCl, and 1 mM DTT. Fractions containing E1 were dialyzed against 20 mM Tris-HCl at pH 7.5 and 50 mM NaCl overnight. The dialyzed sample was applied to a HiTrap Q column (Cytiva) and eluted with a linear gradient from 50 mM to 500 mM NaCl in 20 mM Tris-HCl at pH 7.5. The eluted sample was applied to a GSTrap column; eluted with 50 mM Tris-HCl at pH 8.0, 50 mM NaCl, and 10 mM glutathione; flash-frozen in liquid nitrogen; and stored at −80°C.

### Nucleosome reconstitution

The nucleosome was reconstituted with human core histones and 147- or 185-bp DNA fragments containing the Widom 601 spacing sequence, as described previously (*52*).

### Electrophoretic analysis of the Bre1-nucleosome complex

To monitor the formation of the Bre1-nucleosome complex, the nucleosome and trimeric Bre1 complex were mixed at different molar ratios and NaCl concentrations in 20 mM HEPES at pH 7.5, 1 mM DTT, and 0.5 mM EDTA. The mixtures were incubated for an hour at 4℃ and then analyzed using electrophoresis on non-denaturing 4% acrylamide gel in 0.5% TBE buffer at 150 V for 40 minutes. The gels were stained with SYBR Gold (Thermo Fisher).

### Ubiquitination assay

For the ubiquitination assay using the trimeric Bre1A-Bre1B-Rad6A complex, 1.2 μM 147- or 185-bp nucleosome, 100 nM E1, 0.5–1.5 μM Bre1 complex, and 36 μM ubiquitin were mixed in 50 mM Tris-HCl at pH 8.0, 50 mM NaCl, 50 mM KCl, 3 mM ATP, 10 mM MgCl_2_, and 1 mM DTT and incubated at 30°C for 90 min. For the ubiquitination assay of the truncated RING^A^-RING^B^ heterodimers, 1 μM 147-bp nucleosome, 100 nM E1, 10 μM RING^A^-RING^B^ heterodimer or its mutants, and 36 μM ubiquitin were mixed in 50 mM Tris-HCl at pH 8.0, 125 mM NaCl, 3 mM ATP, 10 mM MgCl_2_, 1 mM DTT, and incubated at 30°C for 1, 2, or 4 hours. Reactions were stopped by the addition of 3× SDS loading buffer and analyzed using SDS-PAGE. Ubiquitinated H2BK120 was detected using ubiquityl-histone H2B (Lys120) (D11), with XPR Rabbit mAb as the primary antibody (No. 5546; Cell Signaling), goat anti-rabbit IgG-HRP (sc-2004; Santa Cruz Biotechnology) as the secondary antibody, and ECL Prime (Cytiva) as the chemiluminescent reagent. The images were recorded using LAS-3000 mini (Fujifilm) and quantified using Multi Gauge v3.0 (Fujifilm).

### Cryo-EM sample preparation and data acquisition

Th nucleosome (147 bp DNA) and Bre1 complex (Bre11A-Bre1B-Rad6A) were mixed in a 1:4 molar ratio. Final concentrations of 4.5 µM NCP(400 µg/mL dsDNA) and 18 µM BRE1 complex were incubated in buffer (20 mM HEPES at pH 7.5, 30 mM NaCl, 1 mM DTT, and 1 mM EDTA) at 4℃ for an hour. The sample was then diluted three times, and 3 µl of the diluted sample was applied onto Cu-Rh grids (quantifoil) and plunge-frozen in liquid ethane using Vitrobot Mark IV (Thermo Fisher Scientific). The images were obtained with a 300-kV Titan Krios G3i microscope (Thermo Fisher Scientific) equipped with a K3 direct electron detector (Gatan), installed at the University of Tokyo, Japan. The data sets were acquired with the SerialEM software (*53*), with a defocus range of −0.8 to −1.8 µm. The data acquisition statistics are shown in Table 1.

### Cryo-EM data processing

Movie stacks were corrected for drift- and beam-induced motion with Relion v3.1 (*54*) and MotionCor2 (*55*). The resulting images were exported to cryoSPARC v2.15.0 (*56*), and the CTF parameters were estimated using Patch CTF. Particles were automatically picked with a blob picker using 100 micrographs and then cleaned with several rounds of two-dimensional (2D) classification. The resulting 2D templates were used for template-based picking from all micrographs using cryoSPARC. After several rounds of 2D classifications, we conducted multiple rounds of *ab initio* reconstruction and heterorefinement with cryoSPARC. Finally, we conducted non-uniform refinement to obtain density maps of the Bre1-nucleosome complex and free nucleosome. The resolution was estimated on the basis of the gold standard Fourier shell correlation curve (FSC) at a 0.143 criterion.

### Model building and refinement

A model structure of the RING^A^-RING^B^ heterodimer (calculated using AlphaFold2) and the nucleosome structure (PDB 1KX5) were fit into the density using UCSF Chimera (*57*) and then manually modified with Coot (*58*). Maps locally sharpened with the phenix.local_aniso_sharpen module in Phenix v1.20.1-4487 (*59*) were used during the model modification. Real-space refinement was performed using the phenix.real_space_refine module in Phenix against non-sharpened maps.

### Figure preparation

Structural figures were prepared using PyMOL v2.5.0 (https://pymol.org/2/) and ChimeraX v1.3 (*60*).

### Data availability

The cryo-EM density maps and models were deposited in the Electron Microscopy Data Bank (EMD-34274 and EMD-34275) and Protein Data Bank (8GUI, 8GUJ, and 8GUK).

## Supporting information

Electron density map and overall structure of Model I

Interaction between arginine anchors and the acidic patch.

A model structure of the RINGA-RINGB-Rad6A-ubiquitin complex bound to the nucleosome.

## Acknowledgments

We thank Dr. Yusuke Sato (Tottori University), who kindly provided the plasmids encoding human E1 and ubiquitin. This study was supported by a grant from the Japan Society for the Promotion of Science KAKENHI (grant No. 21H05161) to T.S.. The study was partially supported by the Platform Project for Supporting Drug Discovery and Life Science Research (Basis for Supporting Innovative Drug Discovery and Life Science Research), funded by the Japan Agency for Medical Research and Development (grant No. JP20am0101115, support No. 1061).

## Author contributions

T.S. conceived the project. S.O., K.U., C.O., S.K., and T.S. prepared the samples. S.O., K.S., T.N., O.N., and T.S. collected cryo-EM images. S.O., K.S., and T.S. solved the cryo-EM structures, with T.N.’s advice. S.O., K.U., C.O., and T.S. performed the biochemical analyses. S.O., K.U., K.O., and T.S. wrote the manuscript.

---

**Supplementary Table 1.**
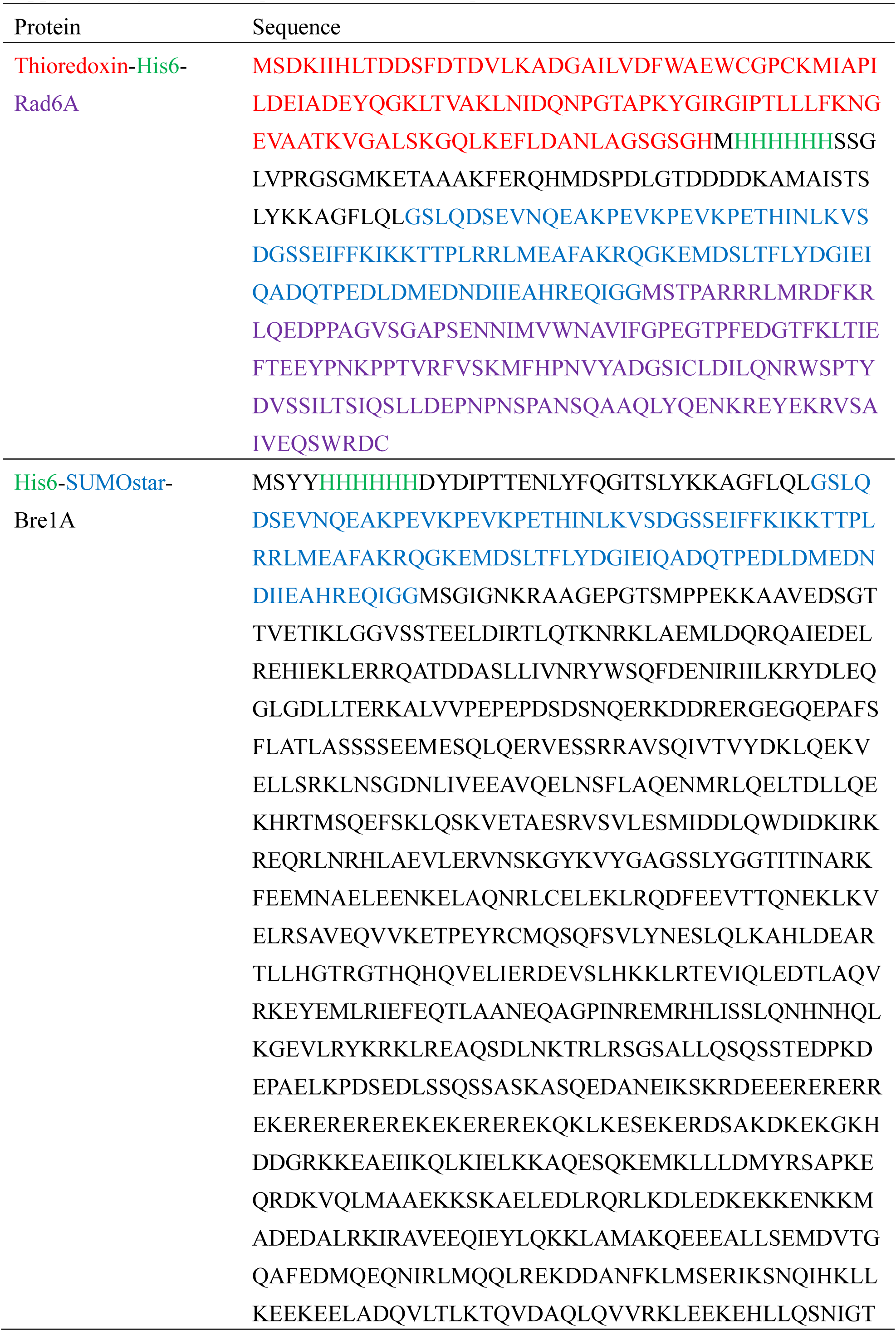
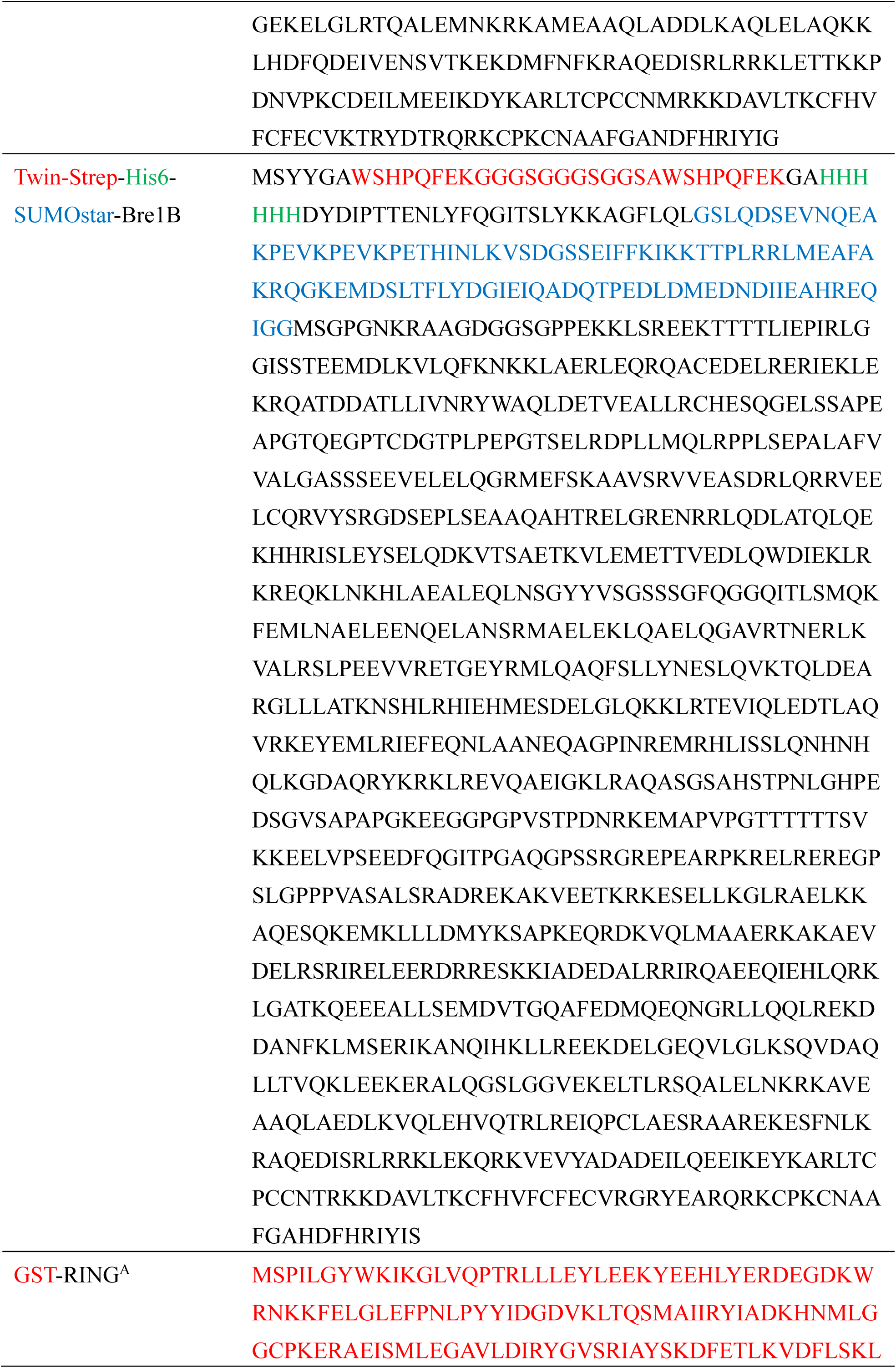
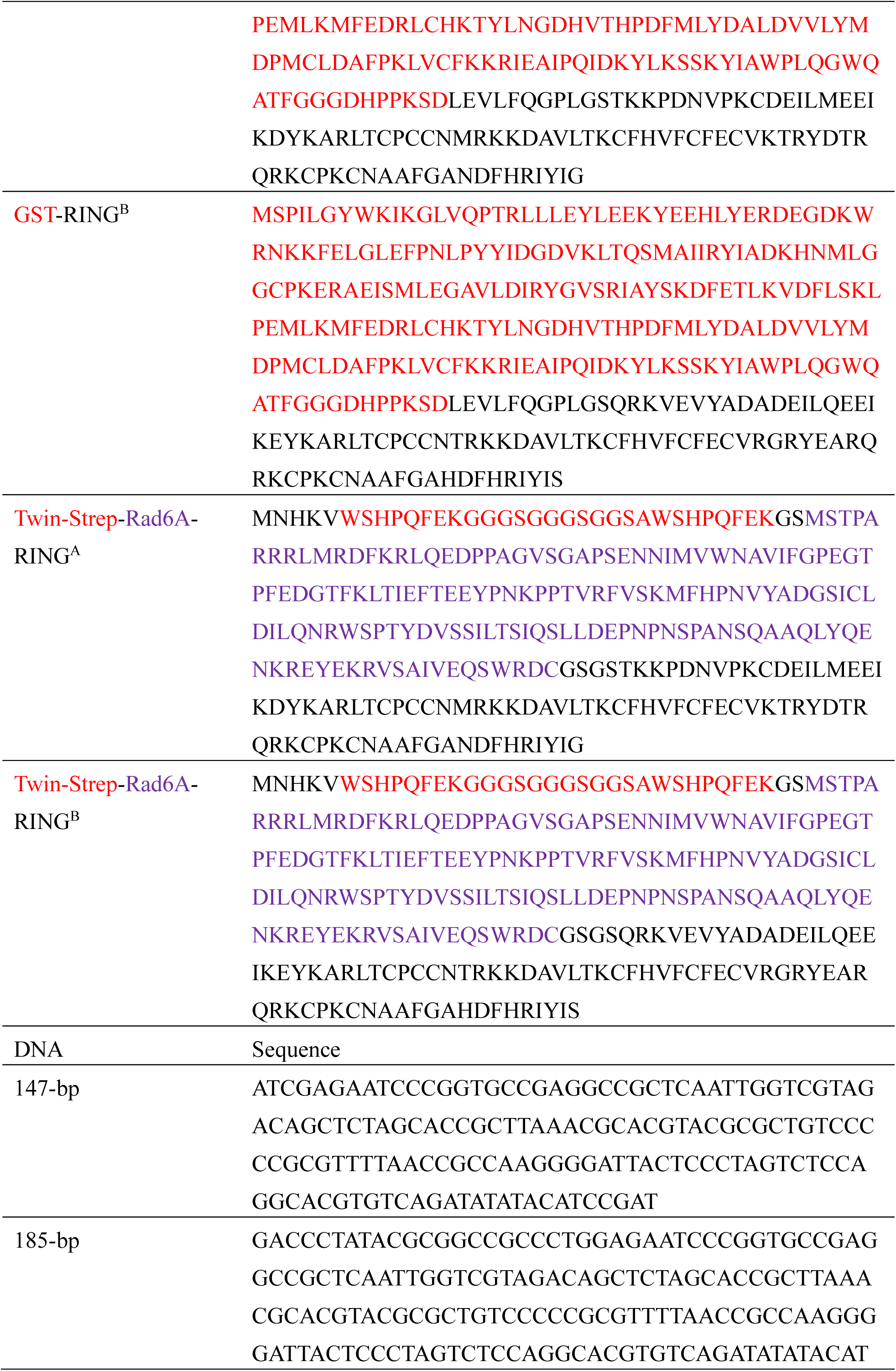
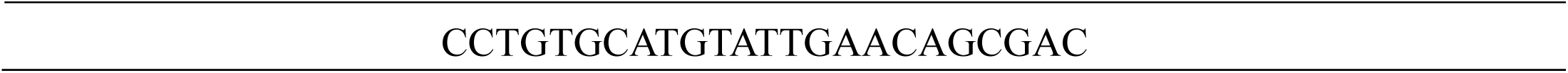
Sequences of recombinant proteins and DNA

**Supplementary Movie 1** Electron density map and overall structure of Model I.

**Supplementary Movie 2** Interaction between arginine anchors and the acidic patch.

**Supplementary Movie 3** A model structure of the RING^A^-RING^B^-Rad6A-ubiquitin complex bound to the nucleosome.

**Extended Data Fig. 1.**
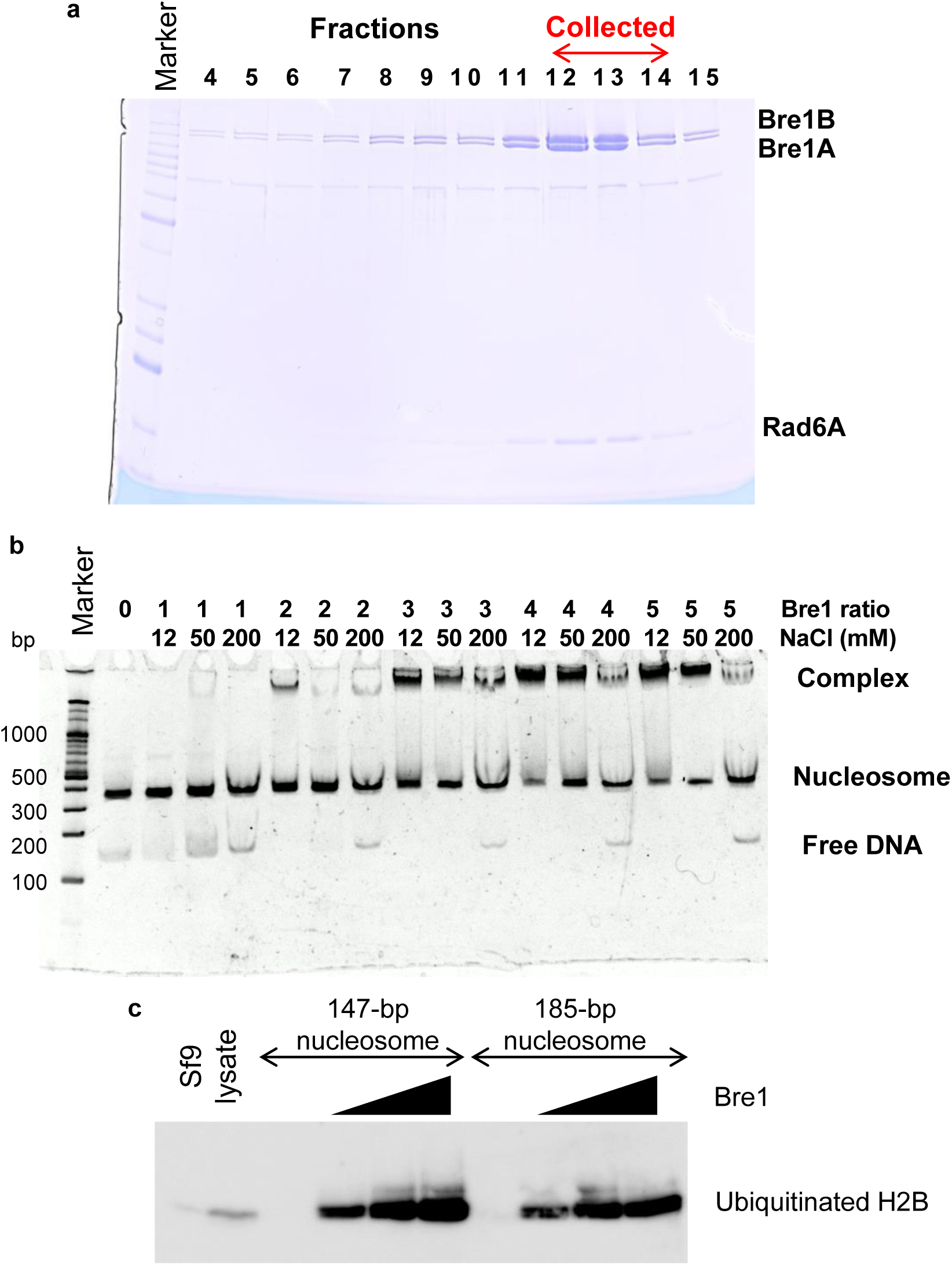
Purification and characterization of the human Bre1 complex. **a**, Purification of the trimeric Bre1A-Bre1B-Rad6A complex using a Superose 6 column. Fractions 12–14 were pooled and used for further study. **b**, Interaction between the trimeric Bre1 complex and the nucleosome with 147-bp DNA analyzed using non-denaturing gel electrophoresis under different Bre1-to-nucleosome molar ratios and NaCl concentrations. **c**, Ubiquitination of nucleosomal H2BK120 by the trimeric Bre1 complex (at 0.5, 1.0, or 1.5 μM) analyzed using western blotting. Sf9 lysate was also loaded as a control.

**Extended Data Fig. 2.**
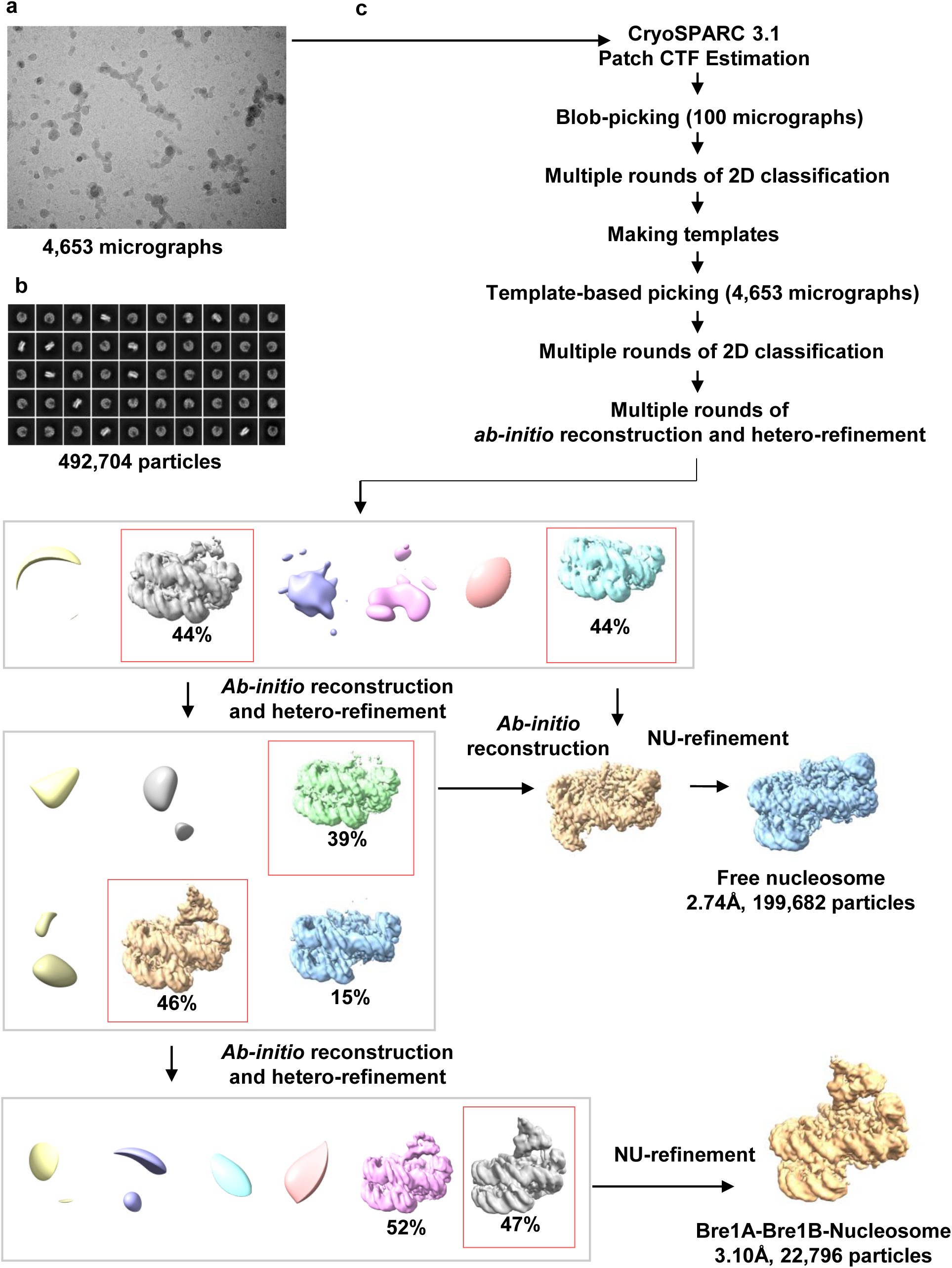
Cryo-EM processing workflow. **a**, Representative motion-corrected micrograph. **b**, Two-dimensional class averages. **c**, Flowchart of data processing.

**Extended Data Fig. 3.**
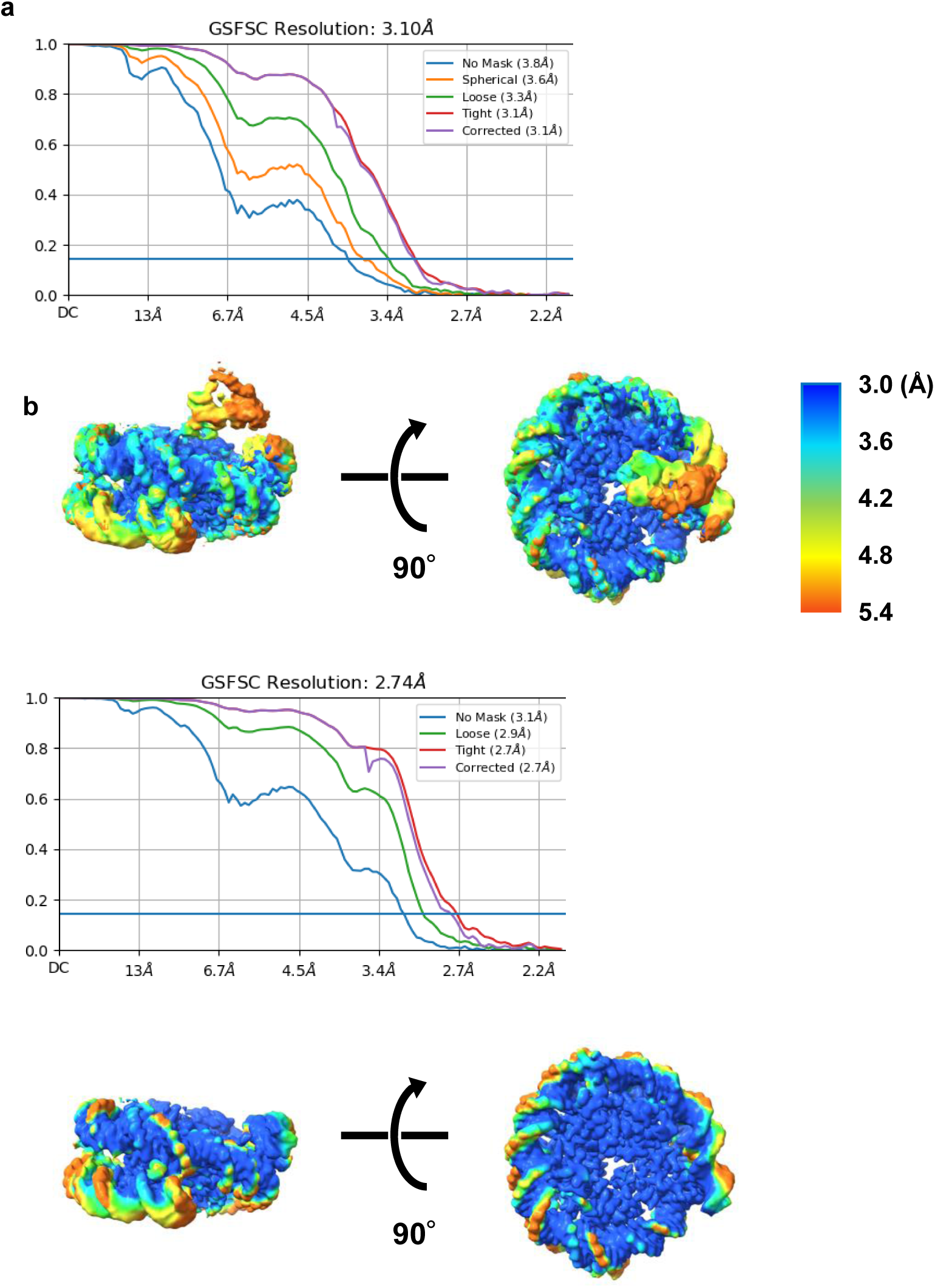
Validation of the cryo-EM maps. **a,** C curve for the Bre1-nucleosome density map calculated using CryoSPARC. **b,** Local resolution map of Bre1-nucleosome. **c**, FSC curve for the free nucleosome density map calculated using CryoSPARC. **d,** Local resolution map of the free nucleosome.

**Extended Data Fig. 4.**
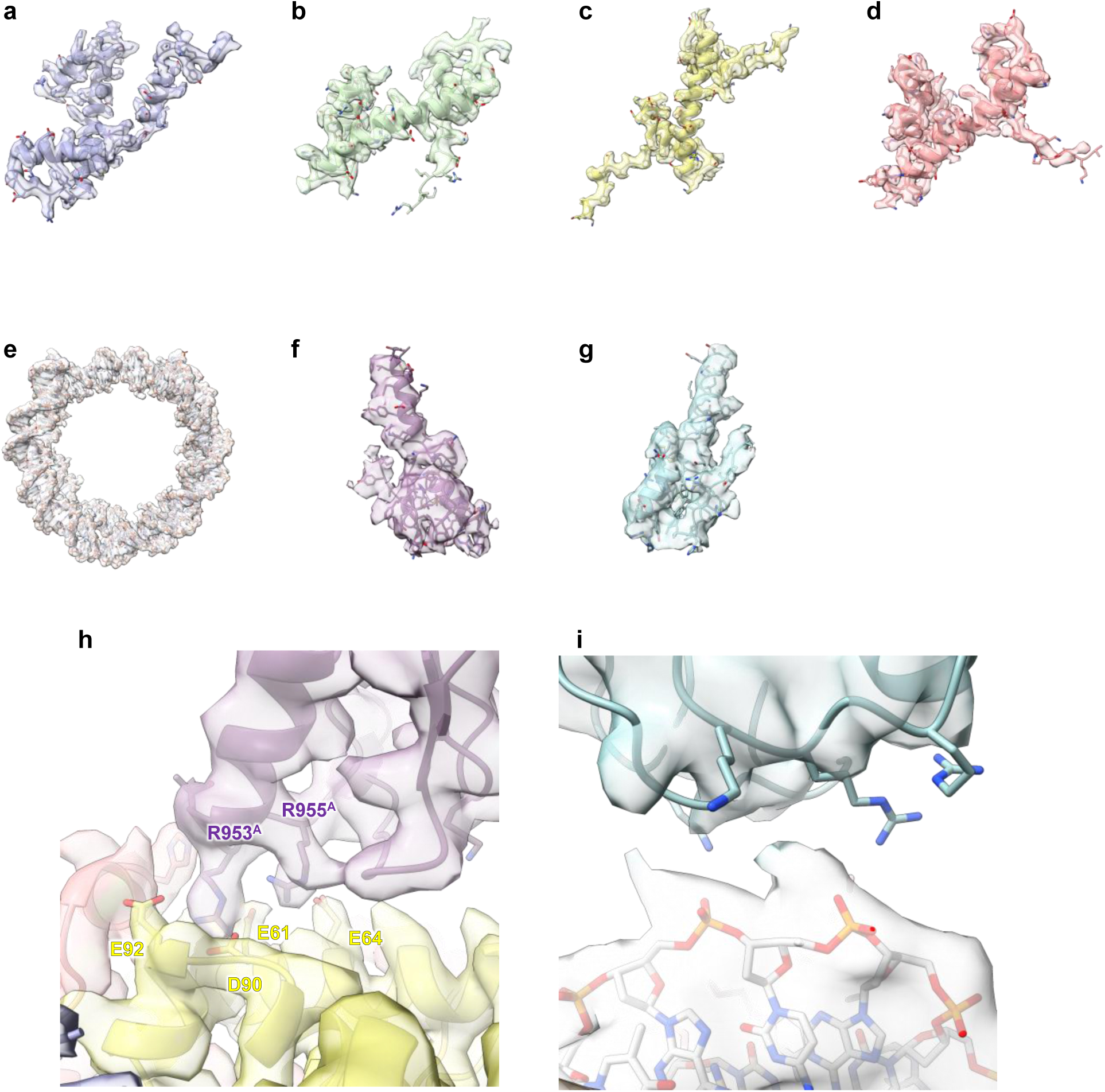
Density map of the Bre1-nucleosome complex. **a,** H3. **b**, H4. **c**, H2A. **d**, H2B. **e**, DNA. **f**, RING domain bound to the acidic patch (modeled here as RING^A^). **g**, RING domain bound to the DNA phosphates (modeled here as RING^B^). **h**, Close-up view near the arginine anchor. **i**, Close-up view of the RING^B^-DNA interface.

**Extended Data Fig. 5.**
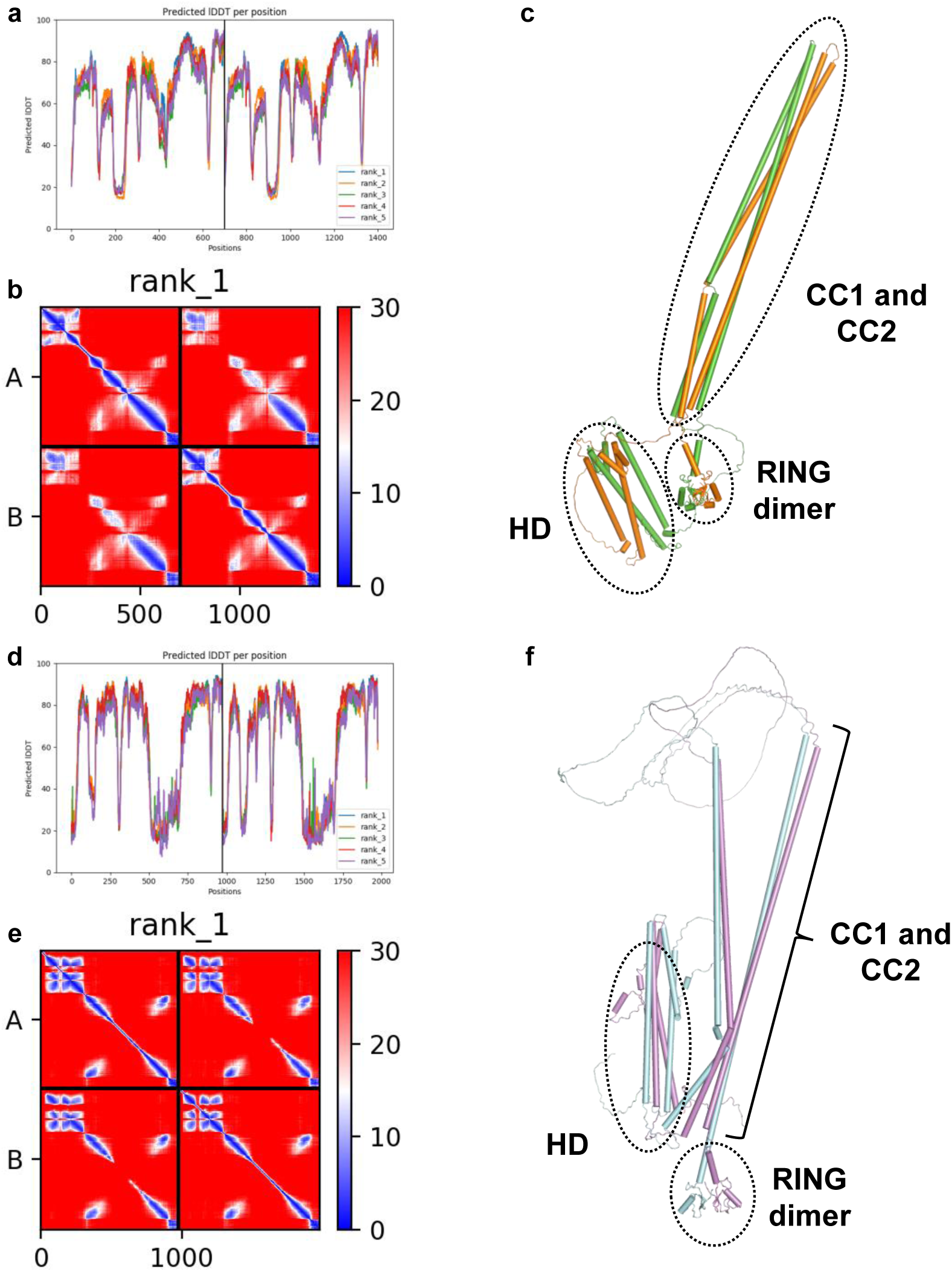
Model structures of yeast (a-c) and human (d-f) Bre1 complexes calculated using AlphaFold. **a, d**, Distribution of the predicted local distance difference test (lDDT) values of five model structures. **b, e**, Distribution of the predicted aligned error (PAE) values (in Å) of the top-ranked model. **c, f**, Overall structure of the top-ranked models. The two yeast Bre1 molecules are colored green and orange.

**Extended Data Fig. 6.**
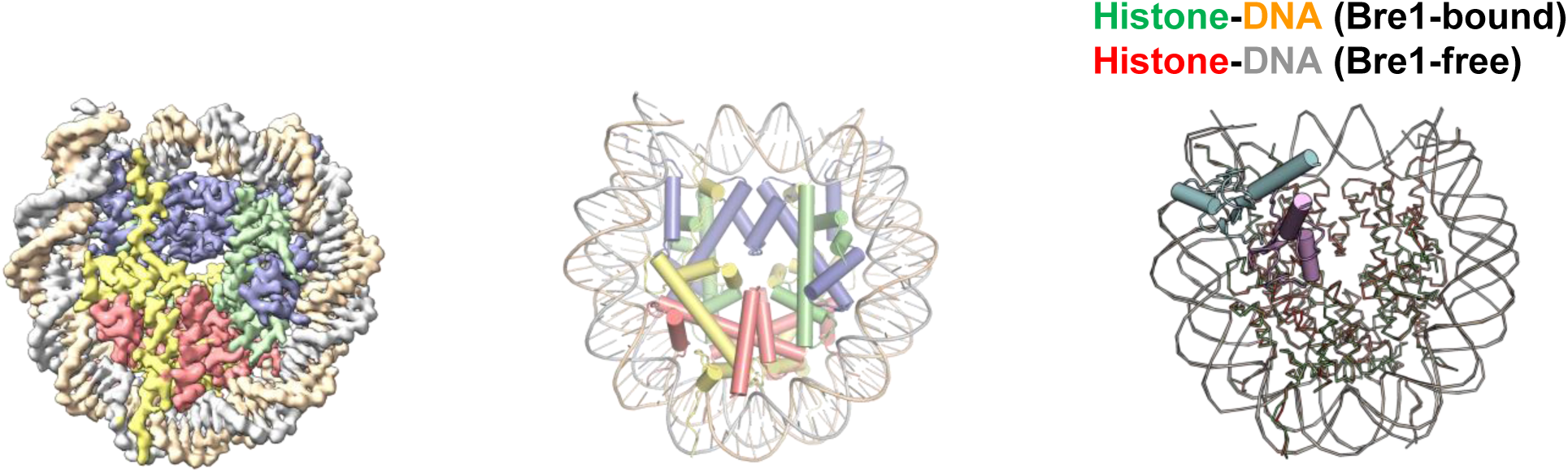
No large conformational change of the nucleosome observed on Bre1 binding. **a**, Density map of the Bre1-free nucleosome at 2.74 Å resolution. **b**, Cartoon representation. **c**, Superposition of the Bre-bound and Bre1-free nucleosome structures determined in this study. RING^A^ and RING^B^ are shown as a cartoon model, whereas histones and DNA are shown as a backbone trace model. Histones and DNA are colored green and orange in the Bre1-bound nucleosomes, and red and gray in the Bre1-free nucleosomes.

**Extended Data Fig. 7.**
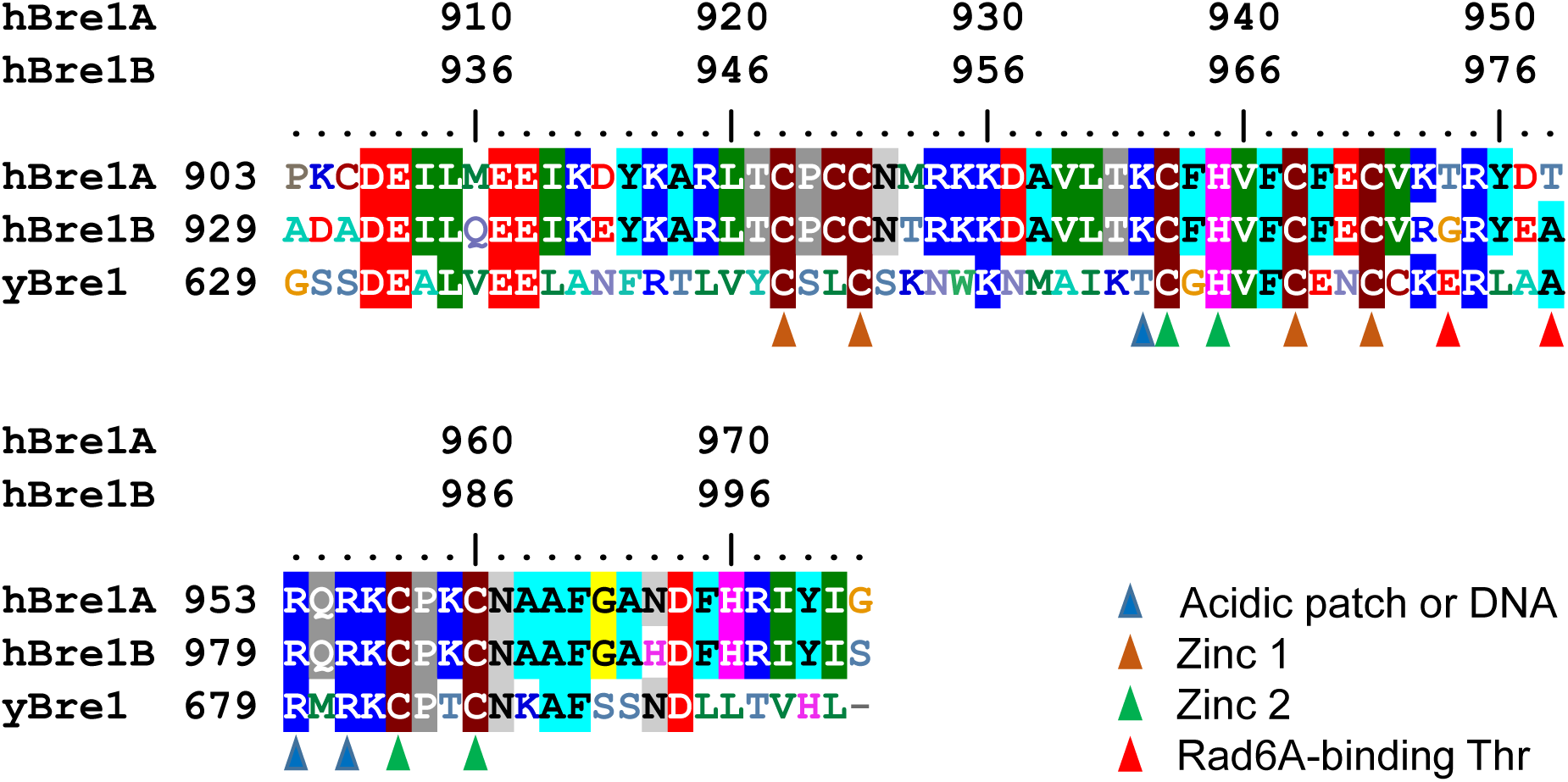
Sequence alignment of the RING domains of yeast Bre1, human Bre1A, and human Bre1B. Residues that contact the acidic patch, DNA phosphates, or zinc ions are indicated. The two threonine residues of Bre1A (T948^A^ and T952^A^) that are possibly involved in Rad6A binding are also indicated.

**Extended Data Fig. 8.**
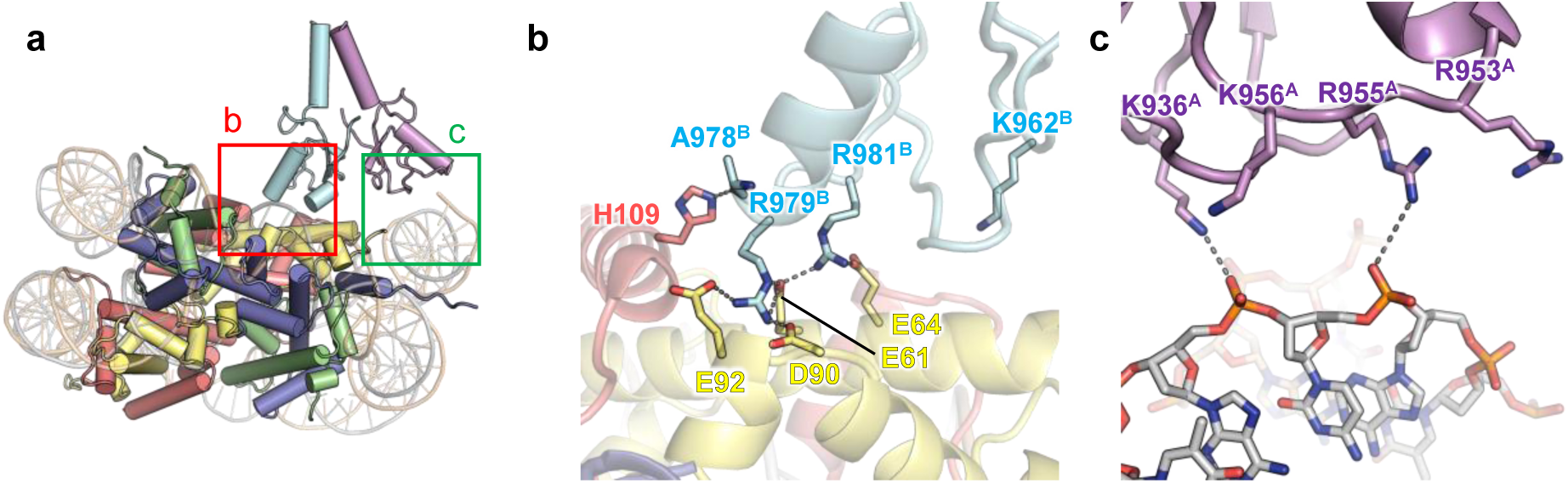
Interactions between the Bre1 complex and the nucleosome in Model II. **a**, Structure of Model II. The magnified regions in b and c are indicated. **b**, Interactions between RING^B^ and the acidic patch. Hydrophilic interactions (salt bridges and hydrogen bonds) are indicated by gray dotted lines. **c**, Basic residues of RING^A^ near the DNA phosphates.

**Extended Data Fig. 9.**
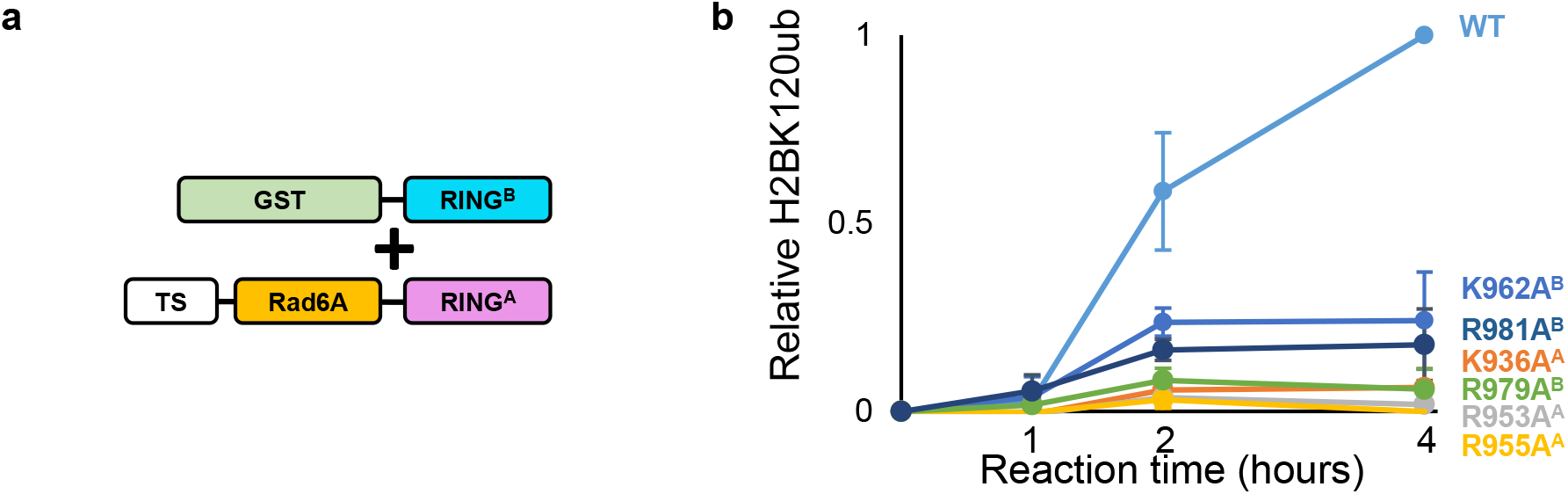
H2BK120 ubiquitination assay using GST-tagged RING^B^ and Twin-Strep-tagged Rad6A-RING^A^ fusion. **a**, Schematic representation of the protein constructs used in the experiment shown in b. TS, Twin-Strep-tag. **b**, H2BK120 ubiquitination assay of the wild type (WT) and mutants possessing substitutions at basic residues. The signals were normalized to the WT activity at 4 hours, and the mean and standard deviation of three independent results are shown.

**Extended Data Fig. 10.**
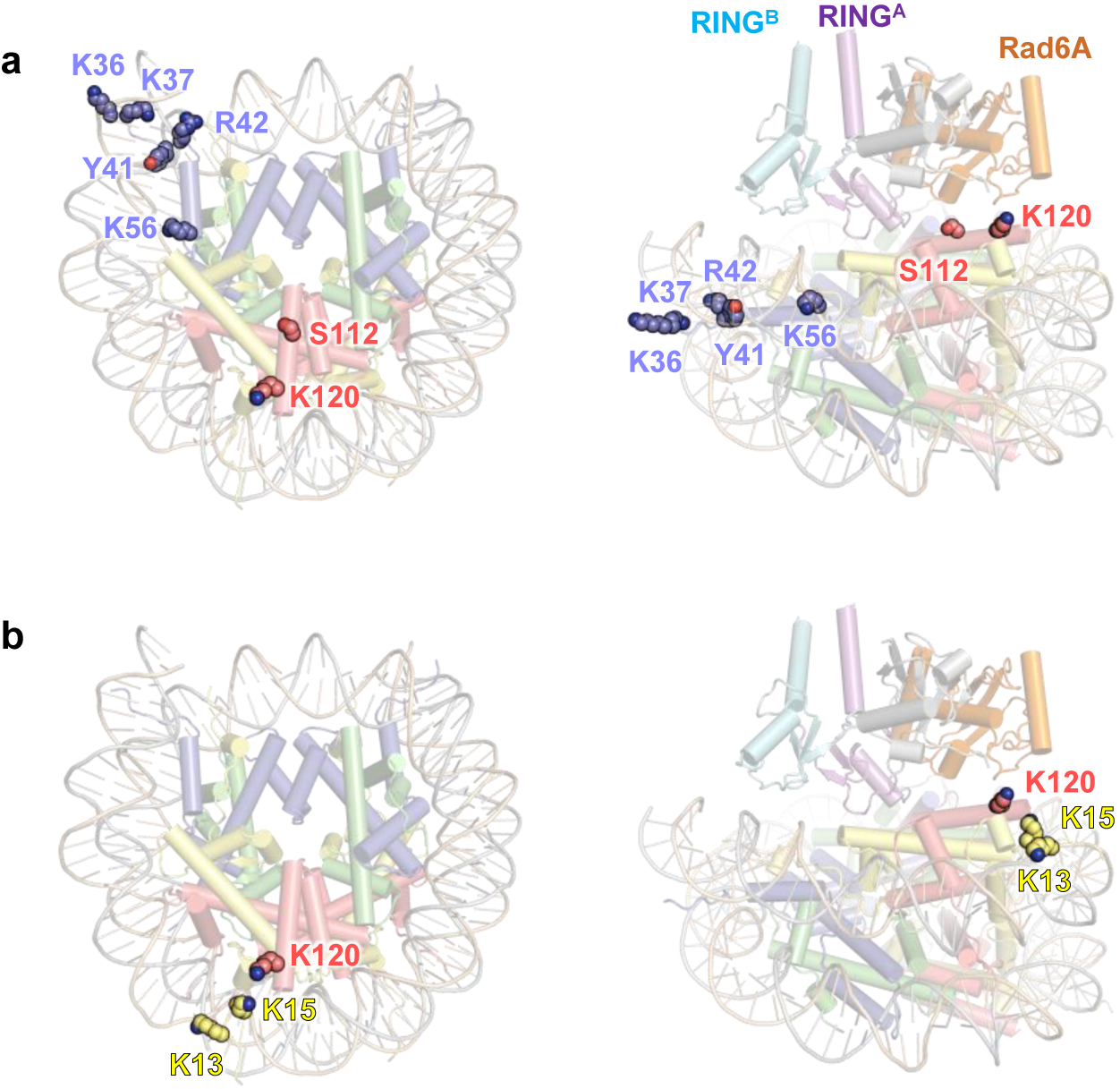
Histone residues whose modifications or substitutions affect the H2BK120 ubiquitination activity by Bre1, as shown by Wojcik *et al*. In the right panels, the model structure of the nucleosome-bound Bre1-Rad6A-ubiquition complex is also shown. (**a**) Residues whose modifications or substitutions (H2BS112GlcNAc, H3K36M, H3K37ac, H3Y41ph, H3R42A, H3K56ac, and H3R42me2a) upregulate Bre1 activity. H2BK120 itself is also shown. (**b**) Residues whose modifications (H2AK13ac, H2AK15ac, H2BK120ac, and H2BK120ub) solely downregulate Bre1 activity.

## References

1. A. J. Bannister, T. Kouzarides, Regulation of chromatin by histone modifications. Cell Res. 21, 381–395 (2011).

2. T. Zhang, S. Cooper, N. Brockdorff, The interplay of histone modifications—writers that read. EMBO Rep. 16, 1467–1481 (2015).

3. R. M. Vaughan, A. Kupai, S. B. Rothbart, Chromatin regulation through ubiquitin and ubiquitin-like histone modifications. Trends Biochem Sci. 46, 258–269 (2021).

4. G. Fuchs, M. Oren, Writing and reading H2B monoubiquitylation. Biochim Biophys Acta. 1839, 694–701 (2014).

5. B. Schuettengruber, H.-M. Bourbon, L. Di Croce, G. Cavalli, Genome regulation by polycomb and trithorax: 70 years and counting. Cell. 171, 34–57 (2017).

6. Z.-W. Sun, C. D. Allis, Ubiquitination of histone H2B regulates H3 methylation and gene silencing in yeast. Nature. 418, 104–108 (2002).

7. J. Dover, J. Schneider, M. A. Tawiah-Boateng, A. Wood, K. Dean, M. Johnston, A. Shilatifard, Methylation of histone H3 by COMPASS requires ubiquitination of histone H2B by Rad6. J Biol Chem. 277, 28368–28371 (2002).

8. S. D. Briggs, T. Xiao, Z.-W. Sun, J. A. Caldwell, J. Shabanowitz, D. F. Hunt, C. D. Allis, B. D. Strahl, Trans-histone regulatory pathway in chromatin. Nature. 418, 498–498 (2002).

9. H. H. Ng, R.-M. Xu, Y. Zhang, K. Struhl, Ubiquitination of histone H2B by Rad6 is required for efficient Dot1-mediated methylation of histone H3 lysine 79. J Biol Chem. 277, 34655–34657 (2002).

10. B. Zhu, Y. Zheng, A.-D. Pham, S. S. Mandal, H. Erdjument-Bromage, P. Tempst, D. Reinberg, Monoubiquitination of human histone H2B: the factors involved and their roles in HOX gene regulation. Mol Cell. 20, 601–611 (2005).

11. J.-S. Lee, A. Shukla, J. Schneider, S. K. Swanson, M. P. Washburn, L. Florens, S. R. Bhaumik, A. Shilatifard, Histone crosstalk between H2B monoubiquitination and H3 methylation mediated by COMPASS. Cell. 131, 1084–1096 (2007).

12. R. K. McGinty, J. Kim, C. Chatterjee, R. G. Roeder, T. W. Muir, Chemically ubiquitylated histone H2B stimulates hDot1L-mediated intranucleosomal methylation. Nature. 453, 812–816 (2008).

13. J. Kim, M. Guermah, R. K. McGinty, J.-S. Lee, Z. Tang, T. A. Milne, A. Shilatifard, T. W. Muir, R. G. Roeder, RAD6-mediated transcription-coupled H2B ubiquitylation directly stimulates H3K4 methylation in human cells. Cell. 137, 459–471 (2009).

14. B. Fierz, C. Chatterjee, R. K. McGinty, M. Bar-Dagan, D. P. Raleigh, T. W. Muir, Histone H2B ubiquitylation disrupts local and higher-order chromatin compaction. Nat Chem Biol. 7, 113–119 (2011ww).

15. R. Pavri, B. Zhu, G. Li, P. Trojer, S. Mandal, A. Shilatifard, D. Reinberg, Histone H2B monoubiquitination functions cooperatively with FACT to regulate elongation by RNA polymerase II. Cell. 125, 703–717 (2006).

16. E. Shema-Yaacoby, M. Nikolov, M. Haj-Yahya, P. Siman, E. Allemand, Y. Yamaguchi, C. Muchardt, H. Urlaub, A. Brik, M. Oren, W. Fischle, systematic identification of proteins binding to chromatin-embedded ubiquitylated H2B reveals recruitment of SWI/SNF to regulate transcription. Cell Rep. 4, 601–608 (2013).

17. A. Wood, N. J. Krogan, J. Dover, J. Schneider, J. Heidt, M. A. Boateng, K. Dean, A. Golshani, Y. Zhang, J. F. Greenblatt, M. Johnston, A. Shilatifard, Bre1, an E3 ubiquitin ligase required for recruitment and substrate selection of Rad6 at a promoter. Mol Cell. 11, 267–274 (2003).

18. W. W. Hwang, S. Venkatasubrahmanyam, A. G. Ianculescu, A. Tong, C. Boone, H. D. Madhani, A Conserved RING finger protein required for histone H2B monoubiquitination and cell size control. Mol Cell. 11, 261–266 (2003).

19. J. Kim, S. B. Hake, R. G. Roeder, The human homolog of yeast BRE1 functions as a transcriptional coactivator through direct activator interactions. Mol Cell. 20, 759–770 (2005).

20. J. Kim, R. G. Roeder, Direct Bre1-Paf1 complex interactions and RING finger-independent Bre1-Rad6 interactions mediate histone H2B ubiquitylation in yeast. J Biol Chem. 284, 20582–20592 (2009).

21. E. Turco, L. D. Gallego, M. Schneider, A. Köhler, Monoubiquitination of histone H2B is intrinsic to the Bre1 RING domain-Rad6 interaction and augmented by a second Rad6-binding site on Bre1. J Biol Chem. 290, 5298–5310 (2015).

22. L. D. Gallego, M. Ghodgaonkar Steger, A. A. Polyansky, T. Schubert, B. Zagrovic, N. Zheng, T. Clausen, F. Herzog, A. Köhler, Structural mechanism for the recognition and ubiquitination of a single nucleosome residue by Rad6–Bre1. Proc Natl Acad Sci U S A. 113, 10553–10558 (2016).

23. R. K. McGinty, S. Tan, Principles of nucleosome recognition by chromatin factors and enzymes. Curr Opin Struct Biol. 71, 16–26 (2021).

24. R. K. McGinty, R. C. Henrici, S. Tan, Crystal structure of the PRC1 ubiquitylation module bound to the nucleosome. Nature. 514, 591–596 (2014).

25. S. R. Witus, A. L. Burrell, D. P. Farrell, J. Kang, M. Wang, J. M. Hansen, A. Pravat, L. M. Tuttle, M. D. Stewart, P. S. Brzovic, C. Chatterjee, W. Zhao, F. DiMaio, J. M. Kollman, R. E. Klevit, BRCA1/BARD1 site-specific ubiquitylation of nucleosomal H2A is directed by BARD1. Nat Struct Mol Biol. 28, 268–277 (2021). https://doi.org/10.1038/s41594-020-00556-4

26. Q. Hu, M. V. Botuyan, D. Zhao, G. Cui, E. Mer, G. Mer, Mechanisms of BRCA1– BARD1 nucleosome recognition and ubiquitylation. Nature. 596, 438–443 (2021).

27. P. Kumar, C. Wolberger, Structure of the yeast Bre1 RING domain. Proteins. 83, 1185– 1190 (2015).

28. M. Foglizzo, A. J. Middleton, C. L. Day, structure and function of the RING domains of RNF20 and RNF40, dimeric E3 ligases that monoubiquitylate histone H2B. J Mol Biol. 428, 4073–4086 (2016).

29. D. J. Marsh, Y. Ma, K.-A. Dickson, Histone monoubiquitination in chromatin remodelling: focus on the histone H2B interactome and cancer. Cancers. 12, 3462 (2020).

30. T. Prenzel, Y. Begus-Nahrmann, F. Kramer, M. Hennion, C. Hsu, T. Gorsler, C. Hintermair, D. Eick, E. Kremmer, M. Simons, T. Beissbarth, S. A. Johnsen, Estrogen-dependent gene transcription in human breast cancer cells relies upon proteasome-dependent monoubiquitination of histone H2B. Cancer Res. 71, 5739–5753 (2011).

31. O. Tarcic, R. Z. Granit, I. S. Pateras, H. Masury, B. Maly, Y. Zwang, Y. Yarden, V. G. Gorgoulis, E. Pikarsky, I. Ben-Porath, M. Oren, RNF20 and histone H2B ubiquitylation exert opposing effects in basal-like versus luminal breast cancer. Cell Death Differ. 24, 694–704 (2017).

32. E. Shema, I. Tirosh, Y. Aylon, J. Huang, C. Ye, N. Moskovits, N. Raver-Shapira, N. Minsky, J. Pirngruber, G. Tarcic, P. Hublarova, L. Moyal, M. Gana-Weisz, Y. Shiloh, Y. Yarden, S. A. Johnsen, B. Vojtesek, S. L. Berger, M. Oren, The histone H2B-specific ubiquitin ligase RNF20/hBRE1 acts as a putative tumor suppressor through selective regulation of gene expression. Genes Dev. 22, 2664–2676 (2008).

33. Y. Urasaki, L. Heath, C. W. Xu, Coupling of glucose deprivation with impaired histone H2B monoubiquitination in tumors. PLoS ONE. 7, e36775 (2012).

34. K.-A. Dickson, A. J. Cole, A. J. Gill, A. Clarkson, G. B. Gard, A. Chou, C. J. Kennedy, B. R. Henderson, Australian Ovarian Cancer Study (AOCS), S. Fereday, N. Traficante, K. Alsop, D. D. Bowtell, A. deFazio, R. Clifton-Bligh, D. J. Marsh, The RING finger domain E3 ubiquitin ligases BRCA1 and the RNF20/RNF40 complex in global loss of the chromatin mark histone H2B monoubiquitination (H2Bub1) in cell line models and primary high-grade serous ovarian cancer. Hum Mol Genet. 25, 5460–5471 (2016).

35. J. Hooda, M. Novak, M. P. Salomon, C. Matsuba, R. I. Ramos, E. MacDuffie, M. Song, M. S. Hirsch, J. Lester, V. Parkash, B. Y. Karlan, M. Oren, D. S. Hoon, R. Drapkin, Early loss of histone H2B monoubiquitylation alters chromatin accessibility and activates key immune pathways that facilitate progression of ovarian cancer. Cancer Res. 79, 760–772 (2019).

36. O. Tarcic, I. S. Pateras, T. Cooks, E. Shema, J. Kanterman, H. Ashkenazi, H. Boocholez, A. Hubert, R. Rotkopf, M. Baniyash, E. Pikarsky, V. G. Gorgoulis, M. Oren, RNF20 links histone H2B ubiquitylation with inflammation and inflammation-associated cancer. Cell Rep. 14, 1462–1476 (2016).

37. K. Zhang, J. Wang, T. R. Tong, X. Wu, R. Nelson, Y.-C. Yuan, T. Reno, Z. Liu, X. Yun, J. Y. Kim, R. Salgia, D. J. Raz, Loss of H2B monoubiquitination is associated with poor-differentiation and enhanced malignancy of lung adenocarcinoma: loss of H2Bub1 enhances lung adenocarcinoma malignancy. Int J Cancer. 141, 766–777 (2017).

38. Z.-J. Wang, Decreased histone H2B monoubiquitination in malignant gastric carcinoma. World J Gastroenterol. 19, 8099 (2013).

39. J. H. Lee, Y. G. Jeon, K.-H. Lee, H. W. Lee, J. Park, H. Jang, M. Kang, H. S. Lee, H. J. Cho, D.-H. Nam, C. Kwak, J. B. Kim, RNF20 suppresses tumorigenesis by inhibiting the SREBP1c-PTTG1 axis in kidney cancer. Mol Cell Biol. 37, e00265–17 (2017).

40. M. A. Hahn, K.-A. Dickson, S. Jackson, A. Clarkson, A. J. Gill, D. J. Marsh, The tumor suppressor CDC73 interacts with the ring finger proteins RNF20 and RNF40 and is required for the maintenance of histone 2B monoubiquitination. Hum Mol Genet. 21, 559–568 (2012).

41. E. Wang, S. Kawaoka, M. Yu, J. Shi, T. Ni, W. Yang, J. Zhu, R. G. Roeder, C. R. Vakoc, Histone H2B ubiquitin ligase RNF20 is required for MLL-rearranged leukemia. Proc Natl Acad Sci. 110, 3901–3906 (2013).

42. J. Jumper, R. Evans, A. Pritzel, T. Green, M. Figurnov, O. Ronneberger, K. Tunyasuvunakool, R. Bates, A. Žídek, A. Potapenko, A. Bridgland, C. Meyer, S. A. A. Kohl, A. J. Ballard, A. Cowie, B. Romera-Paredes, S. Nikolov, R. Jain, J. Adler, T. Back, S. Petersen, D. Reiman, E. Clancy, M. Zielinski, M. Steinegger, M. Pacholska, T. Berghammer, S. Bodenstein, D. Silver, O. Vinyals, A. W. Senior, K. Kavukcuoglu, P. Kohli, D. Hassabis, Highly accurate protein structure prediction with AlphaFold. Nature. 596, 583–589 (2021).

43. R. Evans, M. O’Neill, A. Pritzel, N. Antropova, A. Senior, T. Green, A. Žídek, R. Bates, S. Blackwell, J. Yim, O. Ronneberger, S. Bodenstein, M. Zielinski, A. Bridgland, A. Potapenko, A. Cowie, K. Tunyasuvunakool, R. Jain, E. Clancy, P. Kohli, J. Jumper, D. Hassabis, Protein complex prediction with AlphaFold-Multimer. Bioinformatics. (2021). https://doi.org/10.1101/2021.10.04.463034.

44. M. Mirdita, K. Schütze, Y. Moriwaki, L. Heo, S. Ovchinnikov, M. Steinegger, ColabFold: making protein folding accessible to all. Nat Methods. 19, 679–682 (2022).

45. L. Buetow, D. T. Huang, Structural insights into the catalysis and regulation of E3 ubiquitin ligases. Nat Rev Mol Cell Biol. 17, 626–642 (2016).

46. A. Plechanovová, E. G. Jaffray, M. H. Tatham, J. H. Naismith, R. T. Hay, Structure of a RING E3 ligase and ubiquitin-loaded E2 primed for catalysis. Nature. 489, 115–120 (2012).

47. J. F. de Oliveira, P. F. V. do Prado, S. S. da Costa, M. L. Sforça, C. Canateli, A. T. Ranzani, M. Maschietto, P. S. L. de Oliveira, P. A. Otto, R. E. Klevit, A. C. V. Krepischi, C. Rosenberg, K. G. Franchini, Mechanistic insights revealed by a UBE2A mutation linked to intellectual disability. Nat Chem Biol. 15, 62–70 (2019).

48. M. C. Gambetta, J. Müller, A critical perspective of the diverse roles of O-GlcNAc transferase in chromatin. Chromosoma. 124, 429–442 (2015).

49. R. Fujiki, W. Hashiba, H. Sekine, A. Yokoyama, T. Chikanishi, S. Ito, Y. Imai, J. Kim, H. H. He, K. Igarashi, J. Kanno, F. Ohtake, H. Kitagawa, R. G. Roeder, M. Brown, S. Kato, GlcNAcylation of histone H2B facilitates its monoubiquitination. Nature. 480, 557–560 (2011).

50. F. Wojcik, G. P. Dann, L. Y. Beh, G. T. Debelouchina, R. Hofmann, T. W. Muir, Functional crosstalk between histone H2B ubiquitylation and H2A modifications and variants. Nat Commun. 9, 1394 (2018).

51. J. Fu, L. Liao, K. S. Balaji, C. Wei, J. Kim, J. Peng, Epigenetic modification and a role for the E3 ligase RNF40 in cancer development and metastasis. Oncogene. 40, 465–474 (2021).

52. K. Sato, A. Kumar, K. Hamada, C. Okada, A. Oguni, A. Machiyama, S. Sakuraba, T. Nishizawa, O. Nureki, H. Kono, K. Ogata, T. Sengoku, Structural basis of the regulation of the normal and oncogenic methylation of nucleosomal histone H3 Lys36 by NSD2. Nat Commun. 12, 6605 (2021).

53. D. N. Mastronarde, Automated electron microscope tomography using robust prediction of specimen movements. J Struct Biol. 152, 36–51 (2005).

54. S. H. W. Scheres, RELION: Implementation of a Bayesian approach to cryo-EM structure determination. J Struct Biol. 180, 519–530 (2012).

55. S. Q. Zheng, E. Palovcak, J.-P. Armache, K. A. Verba, Y. Cheng, D. A. Agard, MotionCor2: anisotropic correction of beam-induced motion for improved cryo-electron microscopy. Nat Methods. 14, 331–332 (2017).

56. A. Punjani, J. L. Rubinstein, D. J. Fleet, M. A. Brubaker, cryoSPARC: algorithms for rapid unsupervised cryo-EM structure determination. Nat Methods. 14, 290–296 (2017).

57. E. F. Pettersen, T. D. Goddard, C. C. Huang, G. S. Couch, D. M. Greenblatt, E. C. Meng, T. E. Ferrin, UCSF Chimera—a visualization system for exploratory research and analysis. J Comput Chem. 25, 1605–1612 (2004).

58. P. Emsley, B. Lohkamp, W. G. Scott, K. Cowtan, Features and development of *Coot*. Acta Crystallogr D Biol Crystallogr. 66, 486–501 (2010).

59. P. V. Afonine, B. K. Poon, R. J. Read, O. V. Sobolev, T. C. Terwilliger, A. Urzhumtsev, P. D. Adams, Real-space refinement in *PHENIX* for cryo-EM and crystallography. Acta Crystallogr D Struct Biol. 74, 531–544 (2018).

60. E. F. Pettersen, T. D. Goddard, C. C. Huang, E. C. Meng, G. S. Couch, T. I. Croll, J. H. Morris, T. E. Ferrin, UCSF CHIMERAX : structure visualization for researchers, educators, and developers. Protein Sci. 30, 70–82 (2021).

